# BEHST: genomic set enrichment analysis enhanced through integration of chromatin long-range interactions

**DOI:** 10.1101/168427

**Authors:** Davide Chicco, Haixin Sarah Bi, Jüri Reimand, Michael M. Hoffman

**Author notes:** corresponding author: Michael M. Hoffman. currently at the Peter Munk Cardiac Centre (Davide Chicco). currently at the Massachusetts Institute of Technology (Haixin Sarah Bi).

## Abstract

Transforming data from genome-scale assays into knowledge of affected molecular functions and pathways is a key challenge in biomedical research. Using vocabularies of functional terms and databases annotating genes with these terms, pathway enrichment methods can identify terms enriched in a gene list. With data that can refer to intergenic regions, however, one must first connect the regions to the terms, which are usually annotated only to genes. To make these connections, existing pathway enrichment approaches apply unwarranted assumptions such as annotating non-coding regions with the terms from adjacent genes. We developed a computational method that instead links genomic regions to annotations using data on long-range chromatin interactions. Our method, Biological Enrichment of Hidden Sequence Targets (BEHST), finds Gene Ontology (GO) terms enriched in genomic regions more precisely and accurately than existing methods. We demonstrate BEHST’s ability to retrieve more pertinent and less ambiguous GO terms associated with results of *in vivo* mouse enhancer screens or enhancer RNA assays for multiple tissue types. BEHST will accelerate the discovery of affected pathways mediated through long-range interactions that explain non-coding hits in genome-wide association study (GWAS) or genome editing screens. BEHST is free software with a command-line interface for Linux or macOS and a web interface (http://behst.hoffmanlab.org/).

## Introduction

High-throughput sequencing enables classes of experiments that produce results in the form of genomic regions. Each experiment identifies particular regions like enhancers, binding sites, open chromatin, or transcripts. We often want to summarize the results of these experiments not as regions, but in understandable terms such as especially affected biological processes or molecular functions. When these regions map neatly to individual genes we can use many of the existing *gene set enrichment analysis* (GSEA) or *pathway enrichment analysis* methods [1, 2, 3, 4, 5, 6, 7, 8, 9, 10, 11]. Most of these methods take a gene list from the experiment, tally functional terms (such as Gene Ontology (GO) [12] terms) previously annotated to the genes, and statistically analyze terms with significant enrichment.

Far fewer tools perform pathway enrichment analysis on arbitrary genomic regions without requiring a gene list. The key problem is that, while genes have comprehensive functional term annotations, other genomic regions generally do not. This necessitates somehow connecting non-genic regions to the annotations on genes. GREAT [13] approaches this problem by defining a regulatory domain for each gene that stretches up to either its nearest neighbors on either side or 1 Mbp, whichever is closest. This assumes inherently that non-coding regions relate most strongly to the nearest genes in one dimension. This assumption may prove reasonable for short distances. As the distance from a non-coding region increases, however, it becomes less likely that it interacts directly with the nearest gene. ChIP-Enrich [14] instead uses ENCODE ChIP-seq peak data sets [15] to link genomic regions to a regulated gene, and then uses a logistic regression approach to estimate the probability of each genomic region to be associated to a particular gene set [14]. TAD Pathways [16] selects genome-wide association study (GWAS) signals for a specific human trait or disease, then finds their topologically associating domains (TADs), and finally selects the genes associated to the boundaries of these TADs.

Several assays directly measure which regions of the genome interact, not just along a chromosome, but in three dimensions. These assays include chromosome conformation capture (3C) [17] and Hi-C [18, 19]. Multiple studies show how long-range chromatin interactions between non-coding regions and distal genes prove critical for understanding the phenotypic effects of genetic variants in these regions [20]. For example, non-coding single nucleotide polymorphisms (SNPs) at the *FTO* locus drive obesity through interactions with the distal gene *IRX3* [21, 22]. As another example, an enhancer at the mouse *Lmbr1* locus drives expression of *Shh* necessary for normal limb development [23]. Mutations in this enhancer can result in preaxial polydactyly, a congenital limb malformation [23, 24]. Additionally, a SNP at the *HERC2* locus causes changes in human pigmentation through a long-range chromatin loop with the pigment gene *OCA2* [25]. Long-range interactions with the *PLCB4* promoter identify it as a potential driver of prostate cancer [26].

We introduce a new method, Biological Enrichment of Hidden Sequence Targets (BEHST), to use long-range chromatin interaction information for better genomic set enrichment analysis. BEHST incorporates experimental evidence of these interactions from Hi-C datasets [27]. These datasets include chromatin loops that bring linearly distal regions up to hundreds of kilobases away within spatial proximity.

## Results

BEHST takes advantage of chromatin loops to precisely associate genes to genomic regions, and then generates an enriched list of functional annotations related to those genes. More precisely, BEHST reads a query dataset of genomic regions, and intersects them with chromatin interactions. BEHST identifies gene *cis*-regulatory regions on the other side of the chromatin loop. BEHST then uses g:Profiler [7] to identify enriched functional annotations on these genes. These serve as enriched functional annotations for the initial genomic regions linked to these genes via long-range interactions (Methods, Figure 1, Figure 2).

**Figure 1:**
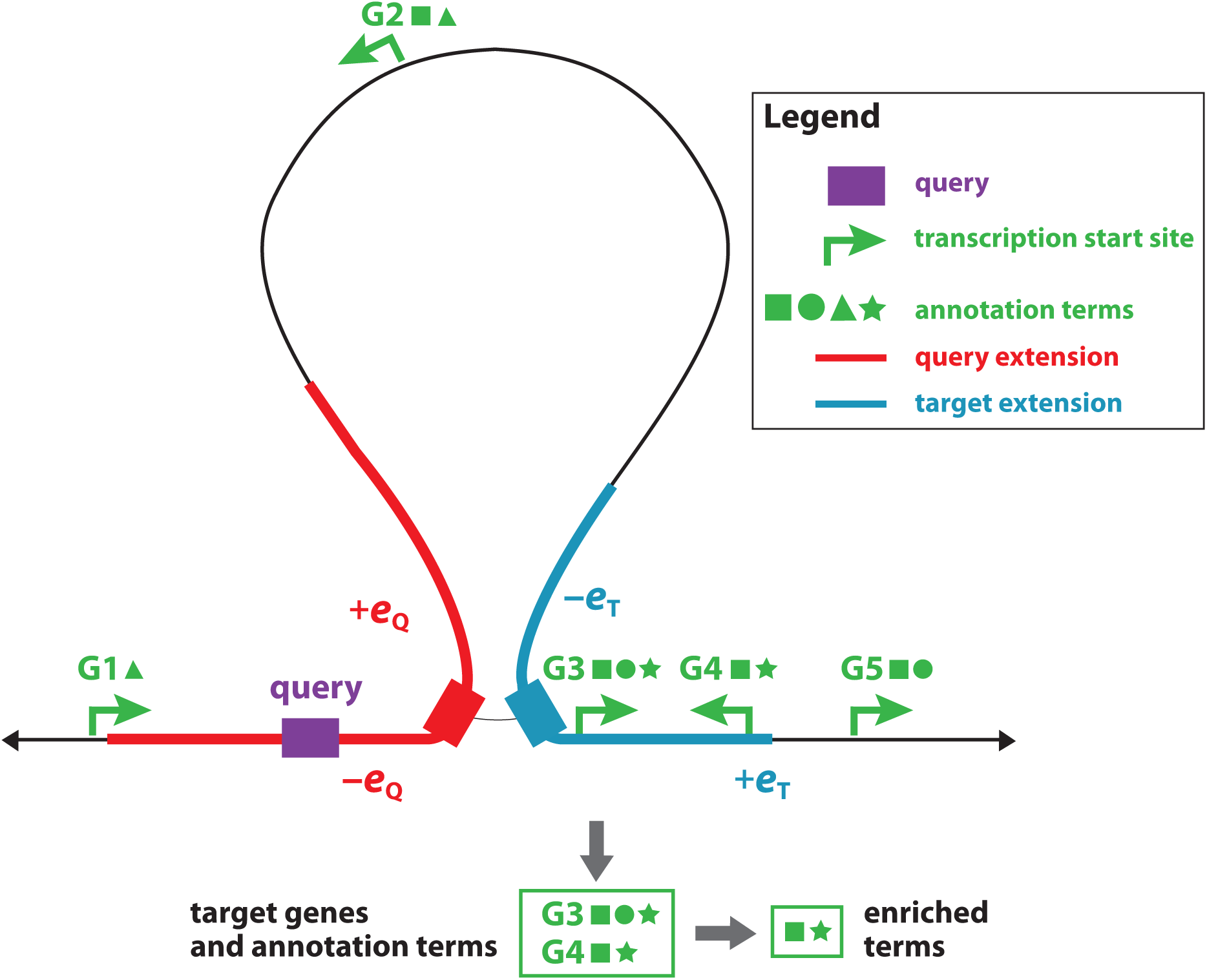
BEHST associates genomic regions with functional annotations on chromatin-loop–linked genes.

BEHST takes a query region (purple), extends the region (red), and intersects it with long-range chromatin interactions (thin black line). On the other side of the interaction, BEHST extends the region (blue), and identifies the gene *cis*-regulatory regions (green arrows) within that extension. Finally, BEHST finds enriched annotations (green symbols) from among the identified genes.

**Figure 2:**
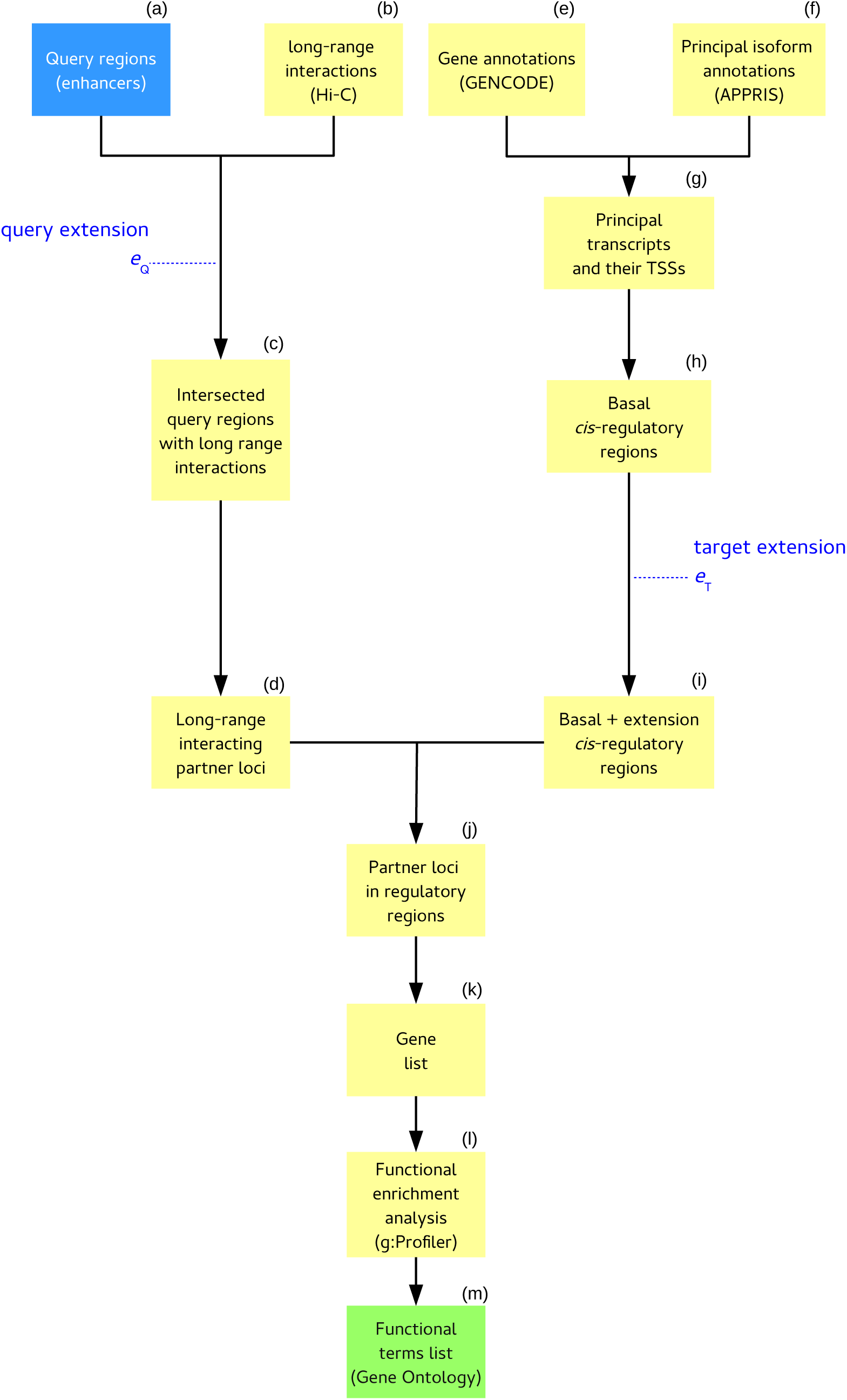
Data flow through BEHST.

**(a)** Blue box: input query regions. **(b, e, f)** Reference datasets shared by multiple BEHST analyses. **(c, d, g–l)** Intermediate data processing steps (Methods).(m) Green box: final resulting enriched GO terms.

### Functional annotations associated with distal enhancers

To examine BEHST’s effectiveness, we used it to identify enriched functional annotations in published enhancers. Both VISTA [28] and FANTOM5 [29] identify sets of enhancers active in particular cell or tissue types. Below, we review how well BEHST enrichment on these datasets recapitulated annotations that one would expect to find for those tissue types.

#### BEHST identifies expected annotations better in VISTA enhancers than in shuffled controls

We applied BEHST to sets of enhancers characterized through transgenic mouse enhancer assays [30], available in the VISTA Enhancer Browser [28]. Each of these 7 datasets included enhancers active in a particular tissue type.

We also compared these VISTA enhancers to two shuffled negative controls. First, we randomly shuffled the enhancers across the whole genome, creating a control input with the same distribution of enhancer size but uncorrelated location. Second, to eliminate effects from moving enhancers between gene-rich and gene-poor regions, we shuffled in a way that preserved distance to the nearest transcription start site (TSS). We did this by identifying the offset between each enhancer and the nearest gene, randomly picking another gene, and moving the enhancer to have the same offset from the new gene (Methods).

BEHST employs two key parameters which control the distance it searches for a chromatin loop from other key regions (Figure 1). The query extension *e*_Q_, defines the distance allowed between a query input region and the nearest chromatin loop. The target extension *e*_T_, defines the distance allowed between the other side of a chromatin loop and the nearest *cis*-regulatory region, where a regulatory region is set as a 6 kbp window (5 kbp upstream and 1 kbp downstream, consistent with GREAT [13]) around the gene’s transcription start site (Methods).

To optimize BEHST’s query and target extension parameters, we performed a grid search. We ran BEHST on each of the 7 VISTA enhancer datasets with 10 different values for each parameter. This entailed running BEHST on 100 different parameter couples for each dataset, or 700 BEHST executions overall. In each of these 700 cases, we recorded the GO term with the most significant *q*-value.

In general, we expected to observe more significant enrichment from the unaltered data than from either of the shuffled controls. BEHST identified more significant annotation enrichment for the unmodified VISTA enhancers than shuffled controls in 6 of 7 tissue types (Table 1 and Figure 3). Heart was the only tissue for which the shuffled controls had more significant enrichment than the experimental enhancers. The heart enhancers led BEHST to retrieve several GO terms related to blood, but we expected this association: since many of annotations in the Gene Ontology relate to blood, they are often present in functional enrichment analyses, even after genomic region shuffles.

**Table 1:** **Mean log_10_ (q-value) of the most significant GO term retrieved by BEHST for various datasets over the whole grid search.** The datasets include the published VISTA enhancers (original dataset) and two kinds of shuffled controls (total genomic random shuffle and TSS distance preserving shuffling). Bold values: most significant q-value between an enhancer set and its shuffled controls.

|  | original<br>dataset | total genomic<br>random shuffle | TSS distance<br>preserving shuffling | number of<br>regions |
| --- | --- | --- | --- | --- |
| eye | **-36.68** | -17.35 | -20.10 | 63 |
| forebrain | **-51.18** | -28.33 | -12.15 | 396 |
| heart | -08.33 | **-14.47** | -06.40 | 96 |
| hindbrain | **-53.01** | -16.78 | -18.09 | 297 |
| limb | **-74.02** | -58.64 | -12.26 | 246 |
| midbrain | **-57.15** | -23.81 | -19.58 | 327 |
| nose | **-47.56** | -15.26 | -17.90 | 78 |

**Figure 3:**
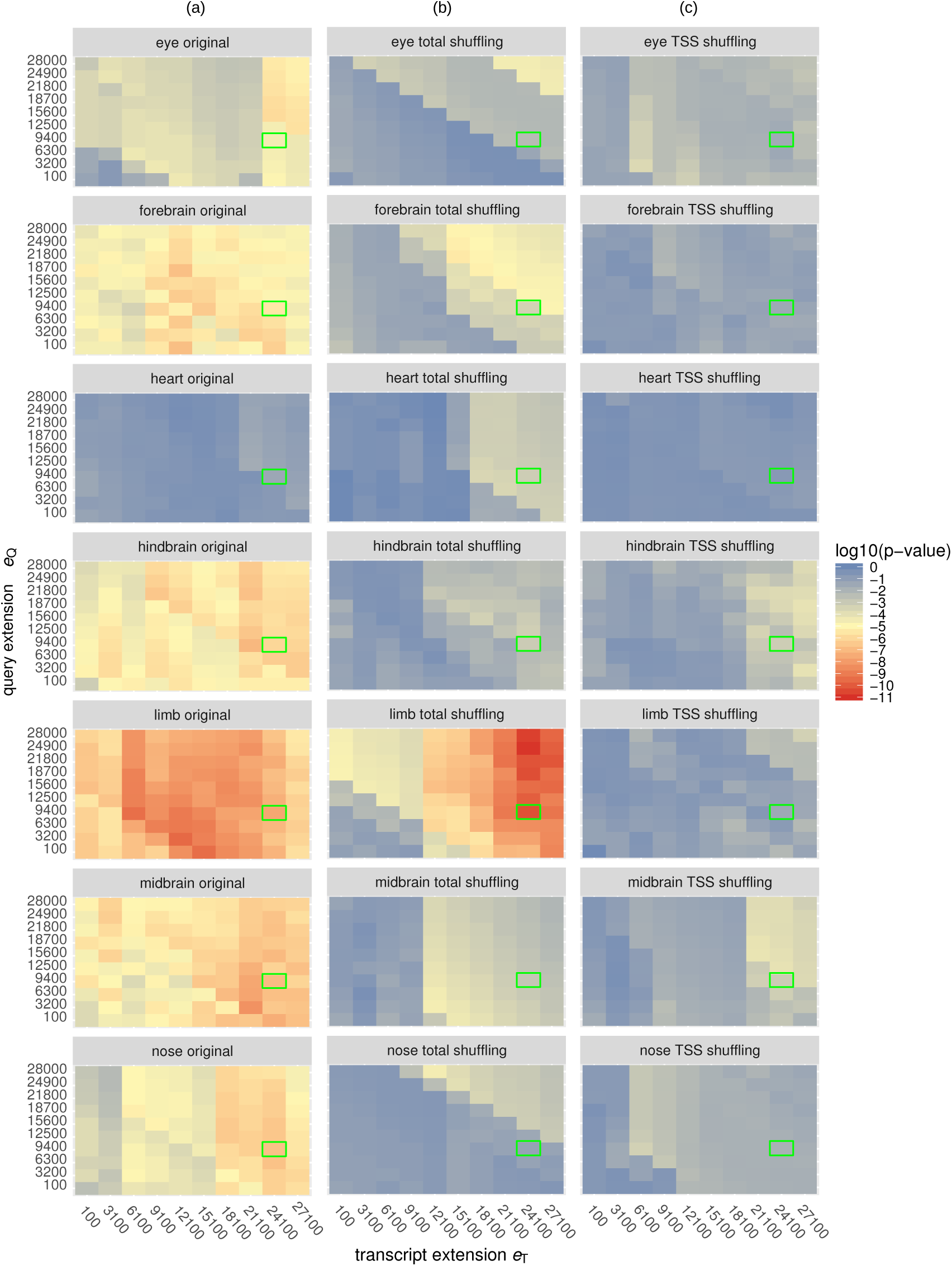
Most significant GO term retrieved by BEHST in a grid search across parameter values.

Each cell represents the log_10_ (q-value) of the most significant GO term found by BEHST for a dataset for a particular parameter couple. Rows within each panel: query extension *e*_Q_, which defines the area around the query region to search for chromatin loops. Columns within each panel: target extension *e*_T_, which defines the area around the distal side of a chromatin loop to search for *cis*-regulatory regions. Green squares highlight the cell containing the optimized parameters (*e*_Q_ = 24,100 bp and *e*_T_ = 9400 bp). Rows of panels: tissue type. Columns of panels: shuffle status: **(a)** Experimental unshuffled VISTA enhancers, **(b)** Control shuffle of the enhancers across the whole genome, **(c)** Control shuffle of the enhancers relative to the nearest transcription start site (TSS).

We used these results to optimize the key extension parameters (Methods). This resulted in optimized values of query extension *e*_Q_ = 24,100 bp, and target extension *e*_T_ = 9400 bp.

#### BEHST can retrieve more specific and more relevant GO terms for VISTA enhancers than existing methods

To examine the enriched GO terms found by BEHST, we focused further on the VISTA limb enhancer and nose enhancer datasets. To aid our evaluation, we manually labeled GO terms with independent association with a particular tissue type. We deemed GO terms with biological relevance to the tissue type as *expected function* (EF), which we analogize to a true positive. We deemed GO terms with biological relevance only to some different tissue type as *unexpected function* (UF), which we analogize to a false positive. Other GO terms, such as those associated with housekeeping functions and many cell types, we do not deem either as expected or unexpected function. Many of these refer to non-specific nucleic acid metabolism processes associated with numerous gene regulation pathways.

##### Limb enhancers

BEHST retrieved multiple expected function terms associated with limb enhancers (Table 2). The most significant term found was “skeletal system development”. BEHST also identified the terms “embryonic limb morphogenesis” and “limb development”. All 81 GO terms found by BEHST (*q* < 0.05) are related to limb, skeleton, embryonic development, or gene regulation.

**Table 2:** **BEHST: VISTA limb**. Most significant 35 GO terms found by BEHST for VISTA limb enhancers. Green rows: terms that refer to limb or skeleton (expected function, EF). Purple rows: terms that refer generally to embryonic development. White rows: terms not specifically related to any tissue. GO: Gene Ontology. BP: biological process. MF: molecular function. *q*: g:Profiler g:SCS q-value [7].

| $q$ | sub-ontology | term ID | EF/UF | term name |
| --- | --- | --- | --- | --- |
| $6.79 \times 10^{-08}$ | BP | GO:0001501 | EF | skeletal system development |
| $2.06 \times 10^{-07}$ | BP | GO:0048598 | | embryonic morphogenesis |
| $2.72 \times 10^{-06}$ | MF | GO:0043565 | | sequence-specific DNA binding |
| $6.27 \times 10^{-06}$ | BP | GO:0009790 | | embryo development |
| $1.94 \times 10^{-05}$ | BP | GO:0007389 | | pattern specification process |
| $3.26 \times 10^{-05}$ | BP | GO:0003002 | | regionalization |
| $3.60 \times 10^{-05}$ | BP | GO:0048562 | | embryonic organ morphogenesis |
| $1.02 \times 10^{-04}$ | BP | GO:0048568 | | embryonic organ development |
| $2.03 \times 10^{-04}$ | BP | GO:0035108 | EF | limb morphogenesis |
| $2.03 \times 10^{-04}$ | BP | GO:0035107 | EF | appendage morphogenesis |
| $2.81 \times 10^{-04}$ | BP | GO:0009059 | | macromolecule biosynthetic process |
| $3.15 \times 10^{-04}$ | BP | GO:0031326 | | regulation of cellular biosynthetic process |
| $3.45 \times 10^{-04}$ | BP | GO:0019438 | | aromatic compound biosynthetic process |
| $3.64 \times 10^{-04}$ | BP | GO:1903506 | | regulation of nucleic acid-templated transcription |
| $4.29 \times 10^{-04}$ | BP | GO:2001141 | | regulation of RNA biosynthetic process |
| $4.32 \times 10^{-04}$ | BP | GO:1901576 | | organic substance biosynthetic process |
| $4.44 \times 10^{-04}$ | BP | GO:0009952 | | anterior/posterior pattern specification |
| $4.54 \times 10^{-04}$ | BP | GO:0048706 | EF | embryonic skeletal system development |
| $4.78 \times 10^{-04}$ | BP | GO:0009889 | | regulation of biosynthetic process |
| $4.88 \times 10^{-04}$ | BP | GO:0010556 | | regulation of macromolecule biosynthetic process |
| $5.11 \times 10^{-04}$ | BP | GO:0034654 | | nucleobase-containing compound biosynthetic process |
| $5.36 \times 10^{-04}$ | BP | GO:0097659 | | nucleic acid-templated transcription |
| $5.74 \times 10^{-04}$ | BP | GO:0032774 | | RNA biosynthetic process |
| $6.43 \times 10^{-04}$ | BP | GO:0035113 | | embryonic appendage morphogenesis |
| $6.43 \times 10^{-04}$ | BP | GO:0030326 | EF | embryonic limb morphogenesis |
| $6.52 \times 10^{-04}$ | BP | GO:0009653 | | anatomical structure morphogenesis |
| $7.54 \times 10^{-04}$ | BP | GO:0048736 | | appendage development |
| $7.54 \times 10^{-04}$ | BP | GO:0060173 | EF | limb development |
| $7.85 \times 10^{-04}$ | BP | GO:0009058 | | biosynthetic process |
| $8.14 \times 10^{-04}$ | BP | GO:0006355 | | regulation of transcription, DNA-templated |
| $8.76 \times 10^{-04}$ | BP | GO:0018130 | | heterocycle biosynthetic process |
| $9.39 \times 10^{-04}$ | BP | GO:1901362 | | organic cyclic compound biosynthetic process |
| $9.74 \times 10^{-04}$ | BP | GO:0009887 | | animal organ morphogenesis |

Unlike in BEHST, limb-related terms did not place highly on GREAT’s most significant terms list. GREAT ranks terms related to limb or skeleton in the lowest positions within the significant GO terms retrieved, such as “embryonic limb morphogenesis” (Table 3, green rows). GREAT missed the limb-associated term “skeletal system development” found by BEHST. Additionally, GREAT found several unexpected function GO terms unrelated to limb: “cardiovascular system development”, “heart development”, and “heart morphogenesis” (Table 3, red rows).

**Table 3:** **GREAT: VISTA limb**. The 24 GO terms found by GREAT for VISTA limb enhancers with *p* < 0.05. Green rows: terms that refer to limb or skeleton (expected function, EF). Purple rows: terms that refer generally to embryonic development. White rows: terms not specifically related to any tissue. Red rows: terms apparently unrelated to limb (unexpected function, UF). GO: Gene Ontology. BP: biological process. MF: molecular function. *p*: binomial rank p-value.

| $p$ | sub-ontology | term ID | EF/UF | term name |
| --- | --- | --- | --- | --- |
| $8.21 \times 10^{-14}$ | MF | GO:0003700 | | sequence-specific DNA binding transcription factor activity |
| $1.04 \times 10^{-13}$ | MF | GO:0001071 | | nucleic acid binding transcription factor activity |
| $2.47 \times 10^{-09}$ | BP | GO:0072358 | UF | cardiovascular system development |
| $3.00 \times 10^{-09}$ | BP | GO:0007507 | UF | heart development |
| $1.06 \times 10^{-08}$ | BP | GO:0035108 | EF | limb morphogenesis |
| $1.07 \times 10^{-08}$ | BP | GO:0060173 | EF | limb development |
| $1.21 \times 10^{-08}$ | BP | GO:0045892 | | negative regulation of transcription, DNA-dependent |
| $1.27 \times 10^{-08}$ | BP | GO:0009887 | | organ morphogenesis |
| $1.33 \times 10^{-08}$ | BP | GO:1901191 | | negative regulation of cellular macromolecule biosynthetic process |
| $2.88 \times 10^{-08}$ | BP | GO:0051253 | | negative regulation of RNA metabolic process |
| $3.31 \times 10^{-08}$ | BP | GO:0035295 | | tube development |
| $3.36 \times 10^{-08}$ | BP | GO:0010629 | | negative regulation of gene expression |
| $3.82 \times 10^{-08}$ | BP | GO:0010558 | | negative regulation of macromolecule biosynthetic process |
| $7.97 \times 10^{-08}$ | BP | GO:0003241 | UF | heart morphogenesis |
| $9.81 \times 10^{-08}$ | BP | GO:0048562 | | embryonic organ morphogenesis |
| $1.42 \times 10^{-07}$ | BP | GO:0030326 | EF | embryonic limb morphogenesis |
| $1.80 \times 10^{-07}$ | BP | GO:0060562 | | epithelial tube morphogenesis |
| $2.19 \times 10^{-07}$ | BP | GO:0035239 | | tube morphogenesis |
| $2.31 \times 10^{-07}$ | MF | GO:0043565 | | sequence-specific DNA binding |
| $2.32 \times 10^{-07}$ | BP | GO:0060429 | | epithelium development |
| $4.26 \times 10^{-07}$ | MF | GO:0000981 | | sequence-specific DNA binding RNA polymerase II transcription factor activity |
| $2.64 \times 10^{-07}$ | BP | GO:0048643 | EF | regulation of skeletal muscle tissue development |
| $4.44 \times 10^{-07}$ | BP | GO:0035115 | EF | forelimb morphogenesis |
| $5.40 \times 10^{-07}$ | BP | GO:0048706 | EF | embryonic skeletal system development |

To examine why GREAT found enriched heart-related terms in a limb enhancer dataset and BEHST did not, we compared the gene lists generated by BEHST and GREAT. ChIP-Enrich does not provide a gene list, so we could not examine its results in the same way. BEHST found 184 genes, while GREAT identified 348 genes. The two sets share 45 genes (Figure 4a–c; *p* = 2.2 × 10^*−*46^; Fisher’s exact test). BEHST retrieves fewer genes than GREAT because it uses more stringent gene selection criteria. Consequently, the GO terms found by BEHST contain fewer unexpected functions.

**Figure 4:**
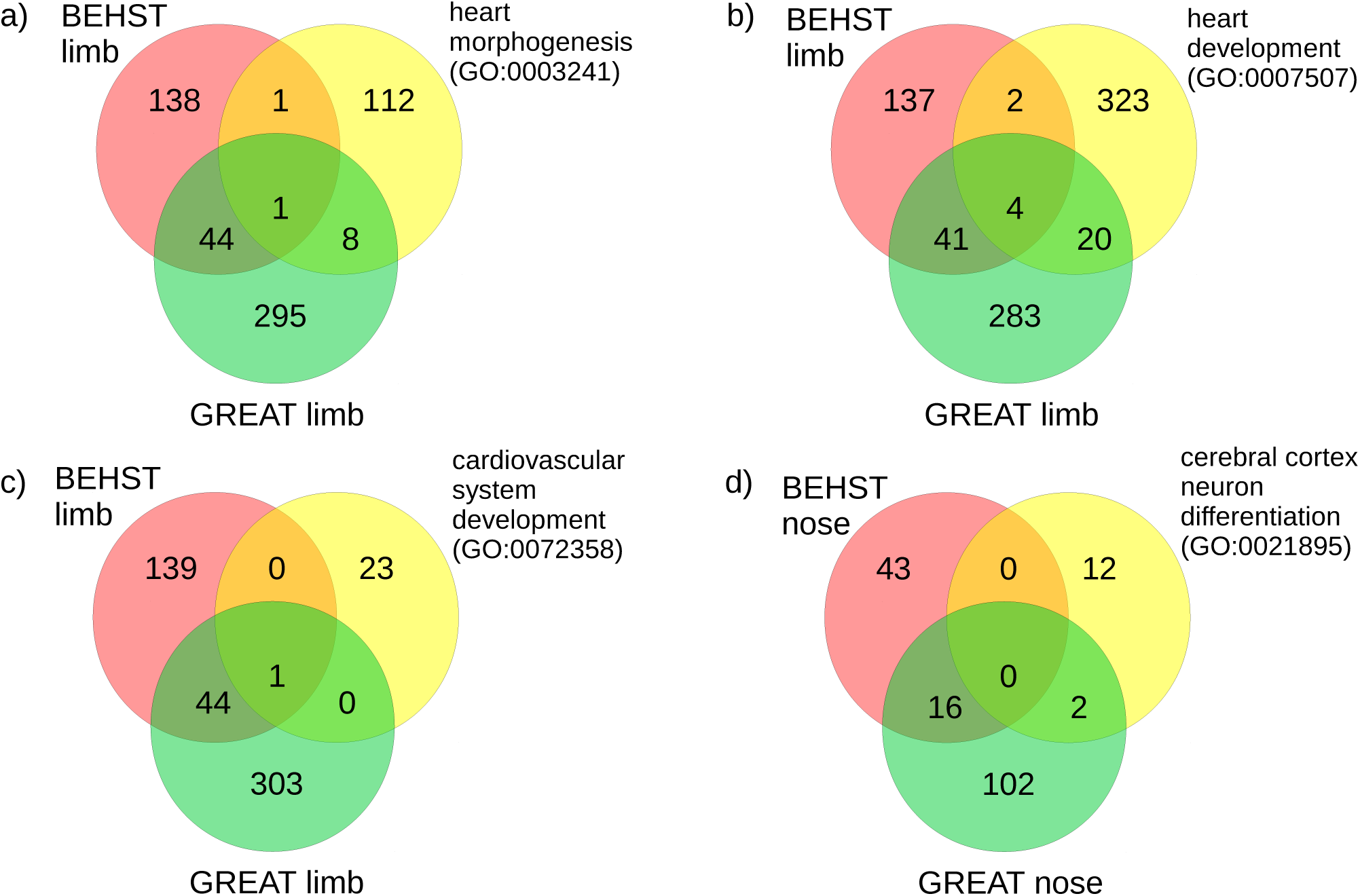
Intersection between gene sets from BEHST, from GREAT, and annotated with particular GO terms.

BEHST and GREAT gene sets for VISTA limb enhancers, intersected with genes annotated with the GO terms **(a)** “heart morphogenesis”, **(b)** “heart development”, and **(c)** “cardiovascular system development”. The BEHST limb set (left red circle) contains 184 genes, and the GREAT limb set contains 348 genes. **(d)** BEHST and GREAT gene sets for VISTA nose enhancers, intersected with genes annotated with the GO term “cerebral cortex neural differentiation”. The BEHST nose set (left red circle) contains 59 genes, and the GREAT nose set (bottom green circle) contains 124 genes.

To further investigate why GREAT, but not BEHST, found enrichment for heart-related terms, we examined how each method performed on individual terms. First, we intersected the GREAT and BEHST limb enhancer gene sets with the set of all genes annotated with “heart morphogenesis” (Figure 4a). The three sets share one common gene, *GJA1* BEHST associates 1 other gene, *TH*, with “heart morphogenesis”, but GREAT associates 8. BEHST’s additional stringency explains why it did not identify an incorrect association with “heart morphogenesis”. We found a similar situation with “heart development”, where BEHST identified 6 genes annotated with this term and GREAT identified 24.

We also intersected the GREAT and BEHST limb enhancer gene sets with the set of all genes annotated with “cardiovascular system development” (Figure 4c). The three sets share one common gene, *FOXB1*, and neither BEHST nor GREAT identify any of the other 23 genes annotated with “cardiovascular system development”.

We also compared BEHST against ChIP-Enrich (Table 4). Like BEHST, ChIP-Enrich identified several expected function GO terms using a conventional approach (Table 4, green rows). Unlike BEHST, it also identified many unexpected function GO terms, clearly unrelated to limb, such as “midbrain-hindbrain boundary development” and “brain development” (Table 4, red rows).

**Table 4:** **ChIP-Enrich: VISTA limb.** Most significant 35 GO terms found by ChIP-Enrich for VISTA limb enhancers. Green rows: terms that refer to limb or skeleton (expected function, EF). Purple rows: terms that refer generally to embryonic development. White rows: terms not specifically related to any tissue. Red rows: terms apparently unrelated to limb (unexpected function, UF). GO: Gene Ontology. BP: biological process. MF: molecular function. CC: cellular component. *p*: binomial rank p-value.

| $p$ | sub-ontology | term ID | EF/UF | term name |
| --- | --- | --- | --- | --- |
| $5.73 \times 10^{-08}$ | MF | GO:0003700 | | sequence-specific DNA binding transcription factor activity |
| $7.51 \times 10^{-08}$ | MF | GO:0001071 | | nucleic acid binding transcription factor activity |
| $2.97 \times 10^{-07}$ | BP | GO:2000272 | | negative regulation of receptor activity |
| $1.72 \times 10^{-06}$ | BP | GO:0035136 | EF | forelimb morphogenesis |
| $2.03 \times 10^{-06}$ | BP | GO:0021532 | EF | neural tube patterning |
| $3.09 \times 10^{-06}$ | BP | GO:0035107 | EF | appendage morphogenesis |
| $4.64 \times 10^{-06}$ | BP | GO:0048538 | | thymus development |
| $1.34 \times 10^{-05}$ | BP | GO:0061099 | | negative regulation of protein tyrosine kinase activity |
| $1.73 \times 10^{-05}$ | BP | GO:0030857 | EF | negative regulation of epithelial cell differentiation |
| $1.73 \times 10^{-05}$ | BP | GO:0030917 | UF | midbrain-hindbrain boundary development |
| $1.77 \times 10^{-05}$ | BP | GO:0030326 | EF | embryonic limb morphogenesis |
| $1.83 \times 10^{-05}$ | BP | GO:0006366 | | transcription from RNA polymerase II promoter |
| $2.34 \times 10^{-05}$ | BP | GO:0050732 | | negative regulation of peptidyl-tyrosine phosphorylation |
| $2.48 \times 10^{-05}$ | BP | GO:0048736 | EF | appendage development |
| $2.81 \times 10^{-05}$ | BP | GO:0021903 | EF | rostrocaudal neural tube patterning |
| $2.96 \times 10^{-05}$ | BP | GO:0009887 | | organ morphogenesis |
| $4.09 \times 10^{-05}$ | BP | GO:0003279 | UF | cardiac septum development |
| $5.04 \times 10^{-05}$ | BP | GO:0060429 | EF | epithelium development |
| $5.20 \times 10^{-05}$ | BP | GO:0050678 | EF | regulation of epithelial cell proliferation |
| $6.16 \times 10^{-05}$ | BP | GO:0050673 | EF | epithelial cell proliferation |
| $9.25 \times 10^{-05}$ | BP | GO:0007420 | UF | brain development |
| $1.18 \times 10^{-04}$ | BP | GO:0006357 | | regulation of transcription from RNA polymerase II promoter |
| $1.73 \times 10^{-04}$ | CC | GO:0000315 | | organellar large ribosomal subunit |
| $1.82 \times 10^{-04}$ | BP | GO:0003205 | UF | cardiac chamber development |
| $1.85 \times 10^{-04}$ | BP | GO:0030855 | EF | epithelial cell differentiation |
| $2.18 \times 10^{-04}$ | BP | GO:0035137 | EF | hindlimb morphogenesis |
| $2.57 \times 10^{-04}$ | BP | GO:0060443 | UF | mammary gland morphogenesis |
| $3.14 \times 10^{-04}$ | BP | GO:0072358 | UF | cardiovascular system development |
| $3.18 \times 10^{-04}$ | BP | GO:0015813 | | L-glutamate transport |
| $3.20 \times 10^{-04}$ | BP | GO:0048863 | | stem cell differentiation |
| $3.48 \times 10^{-04}$ | MF | GO:0043565 | | sequence-specific DNA binding |
| $3.70 \times 10^{-04}$ | BP | GO:0021915 | EF | neural tube development |
| $3.74 \times 10^{-04}$ | BP | GO:0050679 | EF | positive regulation of epithelial cell proliferation |
| $3.95 \times 10^{-04}$ | BP | GO:0000122 | | negative regulation of transcription from RNA polymerase II promoter |
| $3.96 \times 10^{-04}$ | BP | GO:0042733 | | embryonic digit morphogenesis |

##### Nose enhancers

As an additional test case, we applied BEHST to the VISTA nose enhancer dataset. As with limb enhancers, BEHST associated nose enhancers with multiple expected function GO terms such as “skeletal system development”, “embryonic skeletal system development”, and “embryonic skeletal system morphogenesis” (Table 5, green rows). By contrast, GREAT found only one relevant term “cerebral cortex neuron differentiation” at a *p* < 0.05 significance threshold (Table 6). BEHST’s retrieved 59 genes for this dataset, while GREAT retrieved 120 genes. Of these genes, BEHST and GREAT share 16, a significant proportion of annotated genes (*p* = 9.6 *×* 10^*−*24^; Fisher’s exact test). The genes GREAT retrieved with the “cerebral cortex differentiation” term included *ID4* and *ASCL1*, a developmental transcription factor involved in human cerebral cortex neuron differentiation [31].

**Table 5:** **BEHST: VISTA nose.** The 22 GO terms found by BEHST for VISTA nose enhancers with *q* < 0.05. Green rows: terms that refer to limb or skeleton (expected function, EF). Purple rows: terms that refer generally to embryonic development. White rows: terms not specifically related to any tissue. GO: Gene Ontology. BP: biological process. MF: molecular function. *q*: g:Profiler g:SCS q-value [7].

| $q$ | sub-ontology | term ID | EF/UF | term name |
| --- | --- | --- | --- | --- |
| $6.32 \times 10^{-05}$ | MF | GO:0043565 | | sequence-specific DNA binding |
| $2.38 \times 10^{-03}$ | BP | GO:0001501 | EF | skeletal system development |
| $2.81 \times 10^{-03}$ | BP | GO:0048706 | EF | embryonic skeletal system development |
| $3.04 \times 10^{-03}$ | BP | GO:1903506 | | regulation of nucleic acid-templated transcription |
| $3.35 \times 10^{-03}$ | BP | GO:0032774 | | RNA biosynthetic process |
| $3.35 \times 10^{-03}$ | BP | GO:2001141 | | regulation of RNA biosynthetic process |
| $6.18 \times 10^{-03}$ | BP | GO:0051252 | | regulation of RNA metabolic process |
| $6.91 \times 10^{-03}$ | BP | GO:0007389 | | pattern specification process |
| $6.94 \times 10^{-03}$ | BP | GO:0097659 | | nucleic acid-templated transcription |
| $9.39 \times 10^{-03}$ | BP | GO:0019219 | | regulation of nucleobase-containing compound metabolic process |
| $9.67 \times 10^{-03}$ | BP | GO:0010556 | | regulation of macromolecule biosynthetic process |
| $1.03 \times 10^{-02}$ | BP | GO:0003002 | | regionalization |
| $1.06 \times 10^{-02}$ | BP | GO:0006355 | | regulation of transcription, DNA-templated |
| $1.39 \times 10^{-02}$ | BP | GO:0048704 | EF | embryonic skeletal system morphogenesis |
| $2.26 \times 10^{-02}$ | BP | GO:0031326 | | regulation of cellular biosynthetic process |
| $2.37 \times 10^{-02}$ | BP | GO:0006351 | | transcription, DNA-templated |
| $2.79 \times 10^{-02}$ | BP | GO:0010468 | | regulation of gene expression |
| $2.84 \times 10^{-02}$ | BP | GO:0009889 | | regulation of biosynthetic process |
| $2.94 \times 10^{-02}$ | BP | GO:0034654 | | nucleobase-containing compound biosynthetic process |
| $3.81 \times 10^{-02}$ | BP | GO:0051171 | | regulation of nitrogen compound metabolic process |
| $3.95 \times 10^{-02}$ | BP | GO:0018130 | | heterocycle biosynthetic process |
| $4.00 \times 10^{-02}$ | BP | GO:0019438 | | aromatic compound biosynthetic process |

**Table 6:** **GREAT: VISTA nose.** The 1 GO term found by GREAT for VISTA nose enhancers with *p* < 0.05. Red row: term apparently unrelated to nose (unexpected function, UF). GO: Gene Ontology. BP: biological process. *p*: binomial rank p-value.

| $p$ | term ID | sub-ontology | EF/UF | term name |
| --- | --- | --- | --- | --- |
| $3.98 \times 10^{-06}$ | GO:0021895 | BP | UF | cerebral cortex neuron differentiation |

ChIP-Enrich did find the expected function term “nose development” (Table 7, green row). BEHST did not identify any genes with the “nose development” term. Finding this term came at the cost of ChIP-Enrich retrieving many unexpected GO terms (Table 7, red rows) and therefore a loss of specificity.

**Table 7:** **ChIP-Enrich: VISTA nose.** Most significant 35 GO terms found by ChIP-Enrich for VISTA nose enhancers. Green rows: terms that strictly refer to nose (expected functions, EF). Red rows: terms apparently unrelated to nose (unexpected functions, UF). White rows: terms not specifically related to any tissue. GO: Gene Ontology. BP: biological process. MF: molecular function. *p*: binomial rank p-value.

| $p$ | sub-ontology | term ID | EF/UF | term name |
| --- | --- | --- | --- | --- |
| $1.24 \times 10^{-07}$ | BP | GO:0021895 | UF | cerebral cortex neuron differentiation |
| $2.07 \times 10^{-07}$ | BP | GO:0045665 | UF | negative regulation of neuron differentiation |
| $8.76 \times 10^{-07}$ | BP | GO:0030216 | | keratinocyte differentiation |
| $5.90 \times 10^{-06}$ | BP | GO:0035116 | UF | embryonic hindlimb morphogenesis |
| $6.30 \times 10^{-06}$ | BP | GO:0072132 | | mesenchyme morphogenesis |
| $7.27 \times 10^{-06}$ | MF | GO:0043425 | | bHLH transcription factor binding |
| $7.81 \times 10^{-06}$ | BP | GO:0009913 | UF | epidermal cell differentiation |
| $1.48 \times 10^{-05}$ | BP | GO:0009954 | | proximal/distal pattern formation |
| $1.95 \times 10^{-05}$ | BP | GO:0045596 | | negative regulation of cell differentiation |
| $2.24 \times 10^{-05}$ | BP | GO:0045599 | | negative regulation of fat cell differentiation |
| $2.47 \times 10^{-05}$ | BP | GO:0035115 | UF | embryonic forelimb morphogenesis |
| $4.23 \times 10^{-05}$ | BP | GO:0048715 | | negative regulation of oligodendrocyte differentiation |
| $5.11 \times 10^{-05}$ | BP | GO:0048663 | UF | neuron fate commitment |
| $6.45 \times 10^{-05}$ | BP | GO:2000272 | | negative regulation of receptor activity |
| $6.67 \times 10^{-05}$ | MF | GO:0031432 | | titin binding |
| $6.77 \times 10^{-05}$ | BP | GO:0072080 | UF | nephron tubule development |
| $7.17 \times 10^{-05}$ | BP | GO:0006821 | | chloride transport |
| $7.64 \times 10^{-05}$ | BP | GO:0043584 | EF | nose development |
| $8.01 \times 10^{-05}$ | BP | GO:0035137 | UF | hindlimb morphogenesis |
| $8.39 \times 10^{-05}$ | BP | GO:0000288 | | nuclear-transcribed mRNA catabolic process, deadenylation-dependent decay |
| $8.90 \times 10^{-05}$ | BP | GO:0061326 | UF | renal tubule development |
| $1.01 \times 10^{-04}$ | BP | GO:0072079 | UF | nephron tubule formation |
| $1.29 \times 10^{-04}$ | BP | GO:0045616 | | regulation of keratinocyte differentiation |
| $1.33 \times 10^{-04}$ | BP | GO:0045604 | UF | regulation of epidermal cell differentiation |
| $1.41 \times 10^{-04}$ | BP | GO:0007379 | | segment specification |
| $1.73 \times 10^{-04}$ | BP | GO:0045747 | | positive regulation of Notch signaling pathway |
| $1.78 \times 10^{-04}$ | BP | GO:0072078 | UF | nephron tubule morphogenesis |
| $2.09 \times 10^{-04}$ | BP | GO:0035136 | UF | forelimb morphogenesis |
| $2.17 \times 10^{-04}$ | BP | GO:0051093 | | negative regulation of developmental process |
| $2.26 \times 10^{-04}$ | BP | GO:0030857 | | negative regulation of epithelial cell differentiation |
| $2.33 \times 10^{-04}$ | BP | GO:0001709 | | cell fate determination |
| $2.41 \times 10^{-04}$ | BP | GO:0048713 | | regulation of oligodendrocyte differentiation |
| $2.96 \times 10^{-04}$ | BP | GO:0048709 | | oligodendrocyte differentiation |
| $3.20 \times 10^{-04}$ | BP | GO:0021953 | UF | central nervous system neuron differentiation |
| $3.30 \times 10^{-04}$ | BP | GO:0015698 | | inorganic anion transport |

#### BEHST and existing methods retrieve specific and relevant GO terms for FANTOM5 enhancers

To further evaluate the effectiveness of our method, we examined blood enhancers predicted from FANTOM5 cap analysis gene expression (CAGE) data of whole blood [32, 29].

##### Blood enhancers

We examined the FANTOM5 blood enhancers with GREAT, BEHST, and ChIP-Enrich. BEHST found multiple expected function GO terms highly specific for blood, including the top terms “regulation of immune system process”, “immune system process”, and “immune response” (Table 8, green rows). GREAT generated more GO terms in general, and many of these were expected function terms strictly related to blood (Table 9, green rows). ChIP-Enrich even found more expected function GO terms than BEHST and GREAT (Table 10, green rows). None of these methods produced any unexpected function GO terms.

**Table 8:** **BEHST: FANTOM5 blood.** Most significant 35 GO terms found by BEHST for FANTOM5 blood enhancers. Green rows: terms that refer to blood, specifically (expected function, EF). Purple rows: terms that refer generally to blood and immune biology. White rows: terms not specifically related to any tissue. GO: Gene Ontology. BP: biological process. CC: cellular component. *q*: g:Profiler g:SCS q-value [7].

| $q$ | sub-ontology | term ID | EF/UF | term name |
| --- | --- | --- | --- | --- |
| $6.65 \times 10^{-08}$ | BP | GO:0002682 | EF | regulation of immune system process |
| $7.66 \times 10^{-08}$ | BP | GO:0002376 | EF | immune system process |
| $6.77 \times 10^{-07}$ | BP | GO:0006955 | EF | immune response |
| $8.86 \times 10^{-06}$ | BP | GO:0048584 | | positive regulation of response to stimulus |
| $1.75 \times 10^{-05}$ | BP | GO:0002252 | EF | immune effector process |
| $7.53 \times 10^{-05}$ | BP | GO:0002684 | EF | positive regulation of immune system process |
| $1.04 \times 10^{-04}$ | BP | GO:0050776 | EF | regulation of immune response |
| $3.76 \times 10^{-04}$ | BP | GO:0048583 | | regulation of response to stimulus |
| $6.19 \times 10^{-04}$ | BP | GO:0048518 | | positive regulation of biological process |
| $1.09 \times 10^{-03}$ | BP | GO:0001775 | | cell activation |
| $1.22 \times 10^{-03}$ | BP | GO:0002521 | EF | leukocyte differentiation |
| $2.23 \times 10^{-03}$ | BP | GO:0006952 | EF | defense response |
| $3.06 \times 10^{-03}$ | BP | GO:0006950 | | response to stress |
| $3.36 \times 10^{-03}$ | BP | GO:0045321 | EF | leukocyte activation |
| $4.80 \times 10^{-03}$ | BP | GO:1902578 | | single-organism localization |
| $5.86 \times 10^{-03}$ | BP | GO:0002573 | EF | myeloid leukocyte differentiation |
| $6.75 \times 10^{-03}$ | BP | GO:0006909 | | phagocytosis |
| $7.56 \times 10^{-03}$ | CC | GO:0009986 | | cell surface |
| $1.40 \times 10^{-02}$ | BP | GO:0009607 | | response to biotic stimulus |
| $1.46 \times 10^{-02}$ | BP | GO:0002523 | EF | leukocyte migration involved in inflammatory response |
| $1.47 \times 10^{-02}$ | BP | GO:0030097 | EF | hemopoiesis |
| $1.47 \times 10^{-02}$ | BP | GO:0051707 | | response to other organism |
| $1.56 \times 10^{-02}$ | BP | GO:0043207 | | response to external biotic stimulus |
| $1.56 \times 10^{-02}$ | BP | GO:0050778 | | positive regulation of immune response |
| $1.59 \times 10^{-02}$ | BP | GO:0044765 | | single-organism transport |
| $1.62 \times 10^{-02}$ | BP | GO:0006954 | | inflammatory response |
| $1.83 \times 10^{-02}$ | BP | GO:0032496 | EF | response to lipopolysaccharide |
| $2.17 \times 10^{-02}$ | BP | GO:0048522 | | positive regulation of cellular process |
| $2.32 \times 10^{-02}$ | BP | GO:0009605 | | response to external stimulus |
| $2.44 \times 10^{-02}$ | BP | GO:0023014 | | signal transduction by protein phosphorylation |
| $2.56 \times 10^{-02}$ | BP | GO:0070887 | | cellular response to chemical stimulus |
| $2.77 \times 10^{-02}$ | BP | GO:0098602 | | single organism cell adhesion |
| $3.00 \times 10^{-02}$ | BP | GO:0002237 | | response to molecule of bacterial origin |
| $3.10 \times 10^{-02}$ | BP | GO:0046649 | EF | lymphocyte activation |
| $4.89 \times 10^{-02}$ | BP | GO:0007159 | EF | leukocyte cell-cell adhesion |

**Table 9:** **GREAT: FANTOM5 blood.** The 31 GO terms found by GREAT for FANTOM5 blood enhancers with *p* < 0.05. Green rows: terms that refer to blood, specifically (expected functions, EF). Purple rows: terms that refer generally to blood and immune biology. White rows: terms not specifically related to any tissue. GO: Gene Ontology. BP: biological process. MF: molecular function. CC: cellular component. *p*: binomial rank p-value.

| $p$ | sub-ontology | term ID | EF/UF | term name |
| --- | --- | --- | --- | --- |
| $1.44 \times 10^{-71}$ | BP | GO:0002376 | EF | immune system process |
| $3.85 \times 10^{-56}$ | BP | GO:0006952 | EF | defense response |
| $1.09 \times 10^{-50}$ | BP | GO:0006955 | EF | immune response |
| $4.69 \times 10^{-50}$ | BP | GO:0001775 | | cell activation |
| $8.32 \times 10^{-50}$ | BP | GO:0009611 | | response to wounding |
| $1.30 \times 10^{-49}$ | BP | GO:0006954 | | inflammatory response |
| $1.92 \times 10^{-43}$ | BP | GO:0002684 | EF | positive regulation of immune system process |
| $8.85 \times 10^{-43}$ | BP | GO:0002682 | EF | regulation of immune system process |
| $1.80 \times 10^{-39}$ | BP | GO:0045321 | EF | leukocyte activation |
| $2.57 \times 10^{-38}$ | BP | GO:0098542 | | response to other organism |
| $1.38 \times 10^{-37}$ | BP | GO:0009607 | | response to biotic stimulus |
| $6.84 \times 10^{-37}$ | BP | GO:0002407 | | dendritic cell chemotaxis |
| $2.12 \times 10^{-34}$ | BP | GO:0036336 | | dendritic cell migration |
| $2.39 \times 10^{-34}$ | BP | GO:0001817 | | regulation of cytokine production |
| $3.32 \times 10^{-32}$ | BP | GO:0050776 | EF | regulation of immune response |
| $5.61 \times 10^{-32}$ | BP | GO:0046649 | EF | lymphocyte activation |
| $4.06 \times 10^{-28}$ | BP | GO:0002757 | EF | immune response-activating signal transduction |
| $2.54 \times 10^{-27}$ | BP | GO:0050778 | EF | positive regulation of immune response |
| $8.20 \times 10^{-27}$ | BP | GO:0002253 | EF | activation of immune response |
| $4.07 \times 10^{-26}$ | BP | GO:0009617 | | response to bacterium |
| $1.01 \times 10^{-21}$ | MF | GO:0004950 | | chemokine receptor activity |
| $4.97 \times 10^{-16}$ | MF | GO:0004896 | | cytokine receptor activity |
| $8.77 \times 10^{-14}$ | CC | GO:0005944 | | 1-phosphatidylinositol-4-phosphate 3-kinase, class IB complex |
| $5.89 \times 10^{-13}$ | CC | GO:0009897 | EF | external side of plasma membrane |
| $1.95 \times 10^{-9}$ | MF | GO:0043566 | | structure-specific DNA binding |
| $9.64 \times 10^{-8}$ | CC | GO:0001931 | | uropod |
| $3.47 \times 10^{-6}$ | MF | GO:0005178 | | integrin binding |
| $8.19 \times 10^{-6}$ | CC | GO:0070062 | EF | extracellular vesicular exosome |
| $9.89 \times 10^{-6}$ | CC | GO:0065010 | | extracellular membrane-bounded organelle |
| $1.66 \times 10^{-3}$ | CC | GO:0001772 | | immunological synapse |
| $4.59 \times 10^{-3}$ | CC | GO:0031234 | | extrinsic to cytoplasmic side of plasma membrane |

**Table 10:**
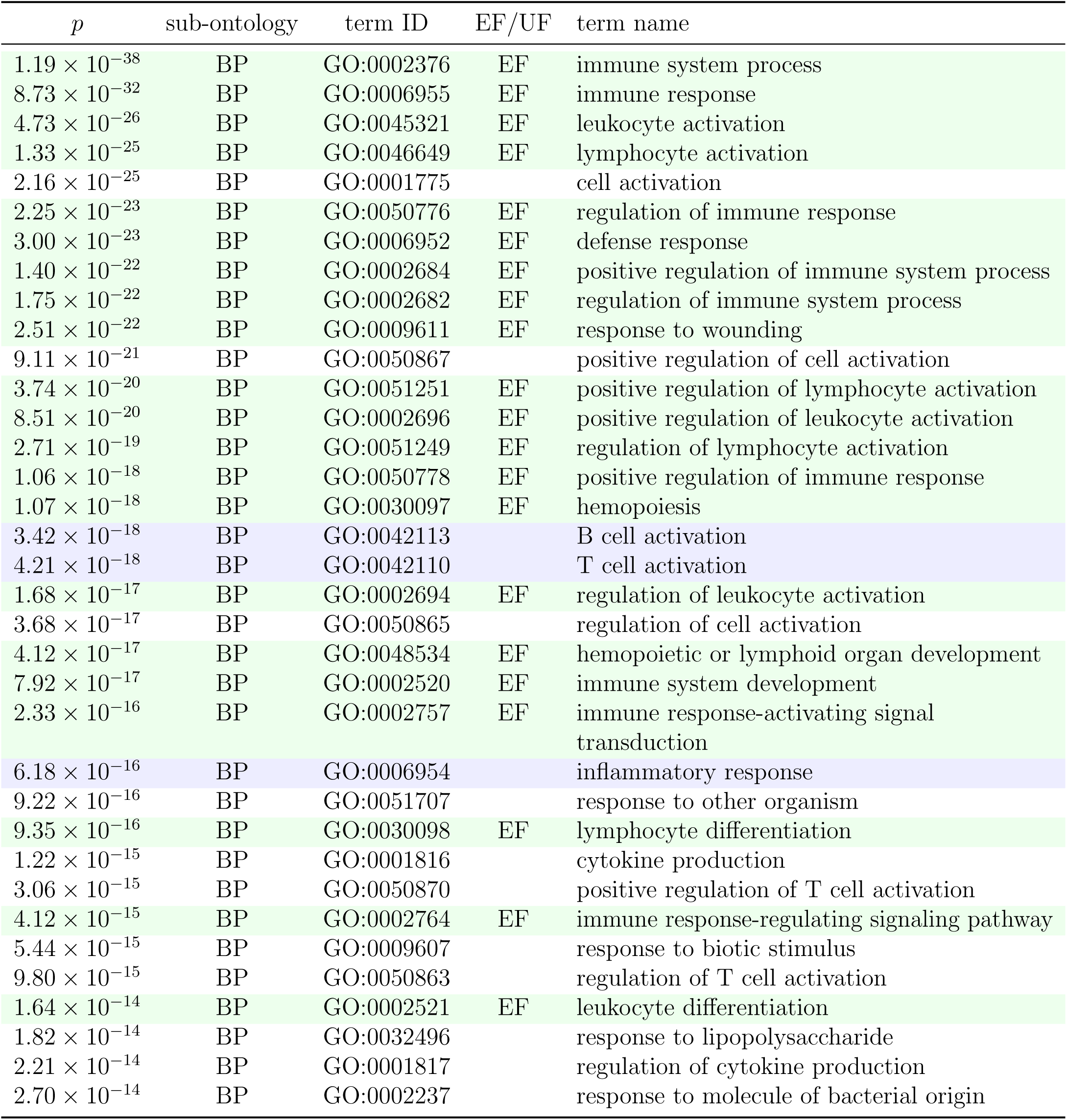
**ChIP-Enrich: FANTOM5 blood.** Most significant 35 GO terms found by ChIP-Enrich for FANTOM5 blood enhancers. Green rows: terms that refer to blood, specifically (expected functions, EF). Purple rows: terms that refer generally to blood and immune biology. White rows: terms that refer to gene regulation. GO: Gene Ontology. BP: biological process. *p*: binomial rank p-value.

#### BEHST’s superior results are robust to different gene set enrichment methods

After identifying target genes, BEHST and GREAT employ different approaches to associate the genes with Gene Ontology terms. GREAT uses a binomial test which explicitly takes into account the variability of gene regulatory domains [13]. BEHST uses g:Profiler [7], which, in turn, employs the g:SCS (set counts and sizes) method [33]. The g:SCS method computes a multiple testing correction for GO term q-values [34]. It considers statistically significant all terms with corrected q-values in the upper fifth percentile.

BEHST, GREAT, and ChIP-Enrich also use different versions of the Gene Ontology Annotation (GOA) [35] database. Here, BEHST used the GOA database of Ensembl 87 (December 2016) [36]. GREAT used a GOA version prior to February 2015. ChIP-Enrich used a GOA version from Bioconductor 2.13 [37], released on October 2013.

Outdated Gene Ontology annotations are a major source of differences in pathway enrichment analyses [1]. We wished to eliminate the possibility that these differences in significance tests or annotation databases drove differences in results between BEHST and GREAT. To do this, we took gene lists produced by GREAT, but did not use GREAT’s binomial test. Instead, we applied g:Profiler to those gene lists. We could not perform a similar analysis using ChIP-Enrich because it does not produce a gene list as output [38].

##### Limb enhancers

First, we ran the hybrid GREAT-g:Profiler analysis on VISTA limb enhancers (Table 12. The hybrid GREAT-g:Profiler analysis found several unexpected function GO terms unrelated to limb, in top positions: “generation of neurons” and “neurogenesis”, among others (Table 12, red rows). This appeared less specific than the enrichment performed by BEHST on the same dataset (Table 2.

**Table 11:** **GREAT-g:Profiler VISTA nose.** The 20 GO terms found by g:Profiler using a GREAT gene list for VISTA nose enhancers with *p* < 0.05. Green rows: terms that strictly refer to nose (expected functions, EF). Purple rows: terms that refer generally to nose biology. Red rows: terms apparently unrelated to nose (unexpected function, UF). GO: Gene Ontology. BP: biological process. CC: cellular component. *q*: g:Profiler g:SCS q-value [7].

| $q$ | sub-ontology | term ID | EF/UF | term name |
| --- | --- | --- | --- | --- |
| $1.25 \times 10^4$ | BP | GO:0022008 | UF | neurogenesis |
| $1.31 \times 10^4$ | BP | GO:0048699 | UF | generation of neurons |
| $3.12 \times 10^4$ | BP | GO:0030182 | UF | neuron differentiation |
| $6.40 \times 10^4$ | BP | GO:0021895 | UF | cerebral cortex neuron differentiation |
| $9.43 \times 10^4$ | BP | GO:0048468 | | cell development |
| $1.63 \times 10^3$ | BP | GO:0021892 | UF | cerebral cortex GABAergic interneuron differentiation |
| $4.35 \times 10^3$ | BP | GO:0021953 | UF | central nervous system neuron differentiation |
| $4.42 \times 10^3$ | BP | GO:0097154 | UF | GABAergic neuron differentiation |
| $5.40 \times 10^3$ | BP | GO:0045595 | UF | regulation of cell differentiation |
| $5.45 \times 10^3$ | BP | GO:0007399 | UF | nervous system development |
| $8.99 \times 10^3$ | BP | GO:0048513 | | animal organ development |
| $9.25 \times 10^3$ | BP | GO:0030154 | | cell differentiation |
| $1.40 \times 10^2$ | BP | GO:0045598 | | regulation of fat cell differentiation |
| $2.29 \times 10^2$ | BP | GO:0021544 | UF | subpallium development |
| $2.35 \times 10^2$ | BP | GO:0048731 | | system development |
| $2.56 \times 10^2$ | BP | GO:0010721 | | negative regulation of cell development |
| $3.29 \times 10^2$ | BP | GO:0048869 | | cellular developmental process |
| $3.98 \times 10^2$ | BP | GO:0030900 | UF | forebrain development |
| $4.04 \times 10^2$ | BP | GO:0035295 | | tube development |
| $4.48 \times 10^2$ | CC | GO:0005634 | | nucleus |

**Table 12:** **GREAT-g:Profiler VISTA limb.** Most significant 35 GO terms found by g:Profiler using a GREAT gene list for VISTA limb enhancers. Green rows: terms that strictly refer to limb (expected functions, EF). Purple rows: terms that refer generally to limb biology. Red rows terms: terms apparently unrelated to limb (unexpected functions, UF). Terms in the white rows refer to gene regulation. GO: Gene Ontology. BP: biological process. MF: molecular function. *q*: g:Profiler g:SCS q-value [7].

| $q$ | sub-ontology | term ID | EF/UF | term name |
| --- | --- | --- | --- | --- |
| $4.69 \times 10^{13}$ | BP | GO:0035295 | | tube development |
| $1.81 \times 10^{11}$ | BP | GO:0009790 | | embryo development |
| $1.92 \times 10^{11}$ | BP | GO:0009887 | | animal organ morphogenesis |
| $2.16 \times 10^{11}$ | BP | GO:0048699 | UF | generation of neurons |
| $3.32 \times 10^{11}$ | BP | GO:0048598 | | embryonic morphogenesis |
| $4.41 \times 10^{11}$ | BP | GO:0022008 | UF | neurogenesis |
| $9.57 \times 10^{11}$ | BP | GO:0007399 | UF | nervous system development |
| $3.18 \times 10^{10}$ | BP | GO:0021537 | | telencephalon development |
| $3.47 \times 10^{10}$ | BP | GO:0030182 | UF | neuron differentiation |
| $4.52 \times 10^{10}$ | BP | GO:0007507 | UF | heart development |
| $1.05 \times 10^{09}$ | BP | GO:0051254 | | positive regulation of RNA metabolic process |
| $1.2 \times 10^{09}$ | BP | GO:0045893 | | positive regulation of transcription,<br>DNA-templated |
| $1.2 \times 10^{09}$ | BP | GO:1903508 | | positive regulation of nucleic acid-templated<br>transcription |
| $1.95 \times 10^{09}$ | BP | GO:0030900 | UF | forebrain development |
| $2.27 \times 10^{09}$ | BP | GO:1902680 | | positive regulation of RNA biosynthetic process |
| $4.53 \times 10^{09}$ | BP | GO:0035239 | | tube morphogenesis |
| $5.99 \times 10^{09}$ | BP | GO:0009653 | EF | anatomical structure morphogenesis |
| $7.37 \times 10^{09}$ | BP | GO:0048562 | | embryonic organ morphogenesis |
| $7.54 \times 10^{09}$ | BP | GO:0045935 | | positive regulation of nucleobase-containing<br>compound metabolic process |
| $7.66 \times 10^{09}$ | BP | GO:0006357 | | regulation of transcription from RNA<br>polymerase II promoter |
| $7.69 \times 10^{09}$ | MF | GO:0003700 | | transcription factor activity, sequence-specific<br>DNA binding |
| $7.96 \times 10^{09}$ | MF | GO:0001071 | | nucleic acid binding transcription factor<br>activity |
| $8.41 \times 10^{09}$ | BP | GO:0003002 | | regionalization |
| $1.63 \times 10^{08}$ | BP | GO:0010628 | | positive regulation of gene expression |
| $3.29 \times 10^{08}$ | BP | GO:0010558 | | negative regulation of macromolecule<br>biosynthetic process |
| $4.27 \times 10^{08}$ | MF | GO:0000981 | | RNA polymerase II transcription factor<br>activity, sequence-specific DNA binding |
| $4.85 \times 10^{08}$ | BP | GO:0051173 | | positive regulation of nitrogen compound<br>metabolic process |
| $5.22 \times 10^{08}$ | BP | GO:0060562 | | epithelial tube morphogenesis |
| $5.35 \times 10^{08}$ | BP | GO:0035107 | EF | appendage morphogenesis |
| $5.35 \times 10^{08}$ | BP | GO:0035108 | EF | limb morphogenesis |
| $5.67 \times 10^{08}$ | BP | GO:0006366 | | transcription from RNA polymerase II<br>promoter |
| $6.36 \times 10^{08}$ | BP | GO:0048513 | | animal organ development |
| $6.49 \times 10^{08}$ | BP | GO:2000113 | | negative regulation of cellular macromolecule<br>biosynthetic process |
| $6.99 \times 10^{08}$ | BP | GO:0043009 | | chordate embryonic development |
| $7.19 \times 10^{08}$ | BP | GO:0006355 | | regulation of transcription, DNA-templated |

##### Nose enhancers

Next, we ran the hybrid GREAT-g:Profiler analysis on VISTA nose enhancers (Table 11). Again, the hybrid analysis found several unexpected function GO terms unrelated to nose in top positions, such as “neurogenesis”, “generation of neurons”, “neuron differentiation”, “cerebral cortex neuron differentiation”, and many others (Table 11, red rows). It also found some GO terms generally related to organ development (for example, “animal organ development), but no expected function GO terms strictly related to nose. This appeared far less specific than BEHST’s enrichment on the same dataset (Table 5).

### Unexpected function retrieval rates in enrichment tests

BEHST’s main goal is providing genomic set enrichment analysis with fewer unexpected function terms than existing tools such as GREAT and ChIP-Enrich. To avoid the strong assumptions inherent in terms such as true positive and false positive, we instead evaluated these methods in terms of the number of expected function terms (EF) and the number of unexpected function terms (UF). In this study, we limited our analysis to the top 35 GO terms with *q* < 0.05 or *p* < 0.05, so the maximum possible values of EF and UF are 35. By analogy to false discovery rate (FDR), we merged EF and UF into a combined measurement called *unexpected function rate* (UFR), where

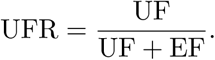

UFR ranges from 0 (best) to 1 (worst).

In addition to measuring performance by comparing UF to those terms specifically designated EF, we also compared it against all of the top 35 GO terms retrieved on a dataset with *q* < 0.05 or *p* < 0.05. The total number of terms found here also includes broadly relevant terms and non-specific terms, such as those pertaining to gene regulation and housekeeping functions. We call this measurement the *total UFR* (tUFR), where

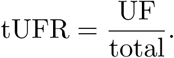

Like UFR, tUFR ranges from 0 (best) to 1 (worst).

To quantitatively summarize the individual comparisons of BEHST and other methods, we computed UFR and tUFR for each comparison (Table 13). In the VISTA tests, BEHST produced UFR and tUFR lower than all the other methods. In FANTOM5 blood enhancers, no method retrieved a UF term, so all methods tied with UFR of 0.00.

**Table 13:** **Summary of expected function (EF) and unexpected function (UF) terms for each dataset examined.** Unexpected function rate UFR = UF*/*(UF+ EF). Total unexpected function rate tUFR = UF*/*total total: number of terms retrieved with *q* < 0.05 or *p* < 0.05, or 35, whichever is smaller.

| dataset | method | total | EF | UF | UFR | tUFR | details |
| --- | --- | --- | --- | --- | --- | --- | --- |
| VISTA limb | **BEHST** | 35 | 6 | 0 | **0.00** | 0.00 | Table 2 |
|  | GREAT | 24 | 6 | 3 | 0.33 | 0.13 | Table 3 |
|  | ChIP-Enrich | 35 | 14 | 5 | 0.26 | 0.14 | Table 4 |
|  | GREAT-g:Profiler | 35 | 3 | 6 | 0.67 | 0.17 | Table 12 |
| VISTA nose | **BEHST** | 22 | 3 | 0 | **0.00** | 0.00 | Table 5 |
|  | GREAT | 1 | 0 | 1 | 1.00 | 1.00 | Table 6 |
|  | ChIP-Enrich | 35 | 1 | 14 | 0.93 | 0.40 | Table 7 |
|  | GREAT-g:Profiler | 20 | 0 | 11 | 1.00 | 0.55 | Table 11 |
| FANTOM5 blood | **BEHST** | 35 | 15 | 0 | **0.00** | 0.00 | Table 8 |
|  | **GREAT** | 31 | 13 | 0 | **0.00** | 0.00 | Table 9 |
|  | **ChIP-Enrich** | 35 | 21 | 0 | **0.00** | 0.00 | Table 10 |

### Semantic similarity of enriched terms

To better understand the differences between BEHST and other methods, we generated semantic similarity analyses of enriched GO terms with REVIGO [39]. For each analysis of a GO term list, REVIGO calculated semantic similarity between every pair of terms in the list. Specifically, we used Resnik similarity [40] to estimate how much information content a pair of terms share [41] in the GO Annotation database [42]. REVIGO removed any redundant terms—terms with a very high semantic similarity with another term. Then, REVIGO clustered enriched terms based upon semantic similarity (Methods).

#### BEHST retrieved EF GO terms as single clusters

In VISTA limb enhancers, BEHST retrieved a cluster of EF GO terms represented by “embryonic skeletal system development” and semantically similar terms (Figure 5a). Thus, BEHST correctly identified a group of similar biological processes, distinct from the rest of the network.

**Figure 5:**
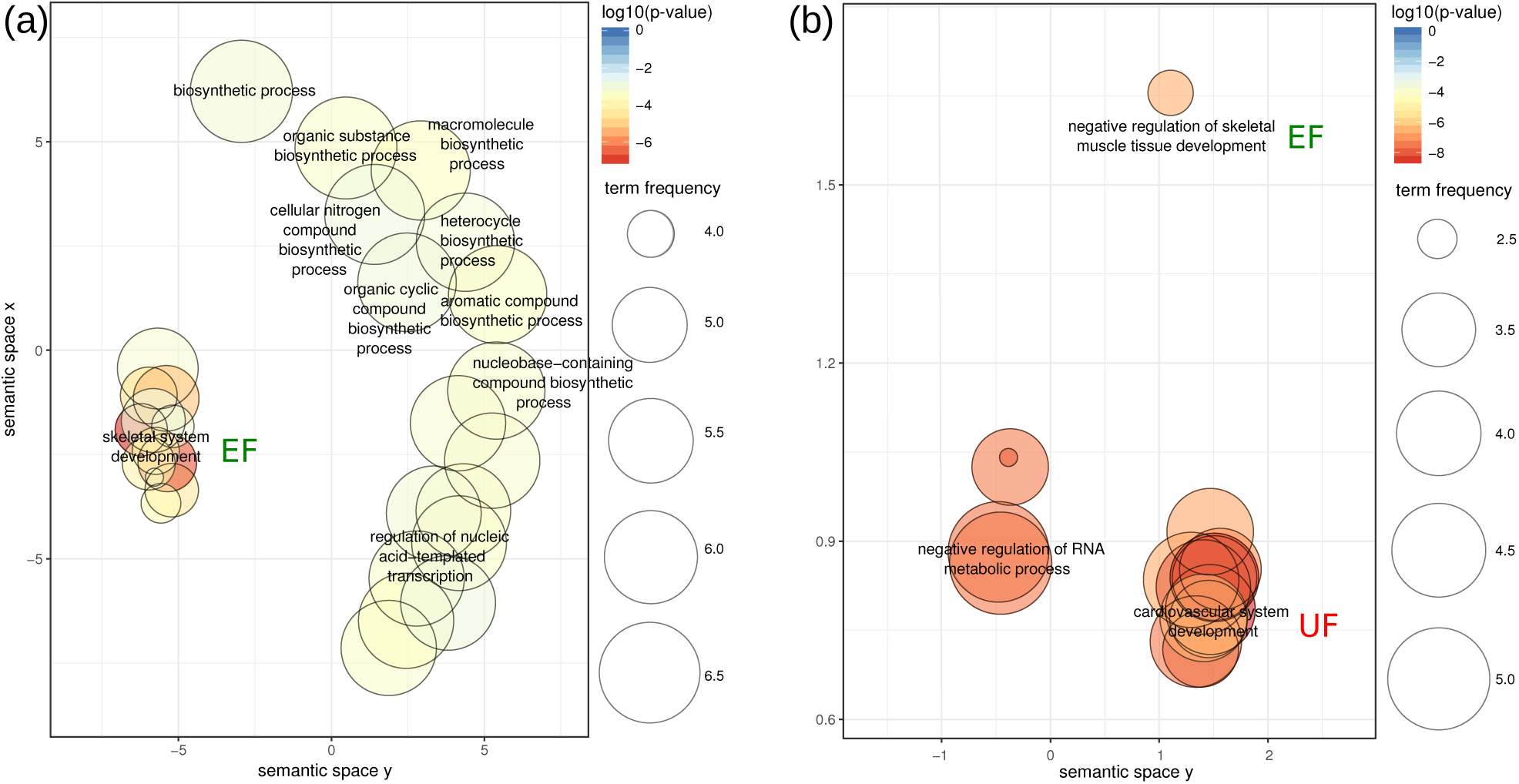
REVIGO [39] scatterplots of GO terms enriched in VISTA limb enhancers.

Enrichment performed by **(a)** BEHST or **(b)** GREAT. Each colored bubble represents a GO term, placed near semantically similar GO terms via multidimensional scaling [43] of a Resnik similarity [40] matrix. Bubble size: background frequency of the term in the GO Annotation database, as a percentage [44]. Bubble color: log_10_ *p*. UF: unexpected function. EF: expected function.

#### GREAT retrieves UF GO terms as accidental errors in clusters of correct GO terms, or as independent UF clusters

For the limb dataset, GREAT found a set of similar GO terms which contains both EFs and UFs (Figure 5b). This cluster shows that GREAT retrieved not only GO terms related to limb, but also included unrelated terms, such as “cardiovascular system development”.

## Discussion

BEHST uses three-dimensional genome organization information instead of adjacency to link arbitrary genomic regions with genes that have annotated GO terms. Using this information, BEHST retrieved more specific and precise GO terms for enriched genomic regions than existing methods. Furthermore, BEHST identified fewer UF GO terms than existing methods, and therefore attained a lower UFR.

We hope to add several extensions to improve BEHST. First, by setting the extension parameters *e*_Q_ and *e*_T_ based on the effective resolution of the long-range interaction data (Table 14, BEHST might provide analyses more tuned to the capabilities of the original experiments. Second, instead of the union of multiple cell types, using chromosome conformation data from only the most relevant cell types could provide more precise enrichment results. Third, adding transcript-centric analysis would allow more use of specific terms annotated to alternative transcripts, where available.

**Table 14:**
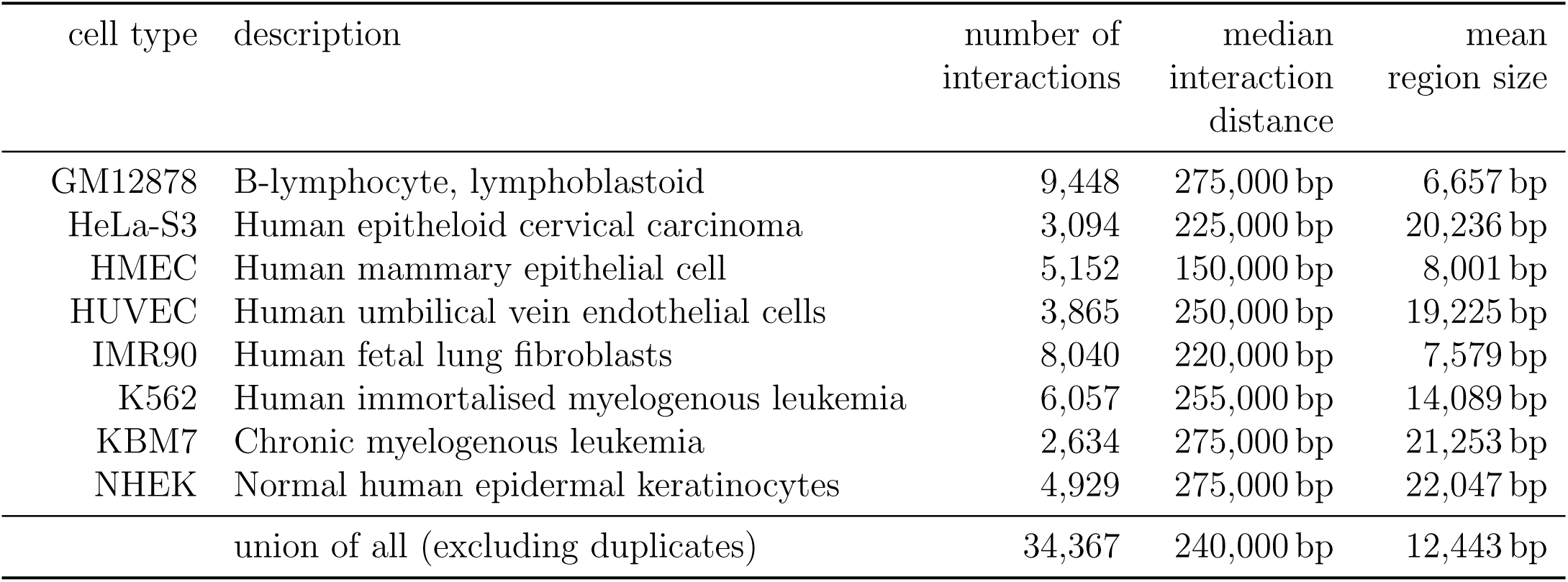
**Summary statistics of the Hi-C datasets used.**

## Methods

### Datasets

In addition to the query data, BEHST employs three reference datasets:

- Long-range interactions: Hi-C data (GEO accession GSE63525 [27]) from a union of eight cell types (Table 14; Table 15);
- Gene annotations: the GENCODE comprehensive gene annotation (version 19 GRCh37.p13 [45]);
- Principal transcript annotations: APPRIS principal isoform annotation (version 2017 01.v20, Species: Human, Assembly Version: GRCh37/hg19, Gene Dataset: Gencode19/Ensembl74 [46]).

**Table 15:**
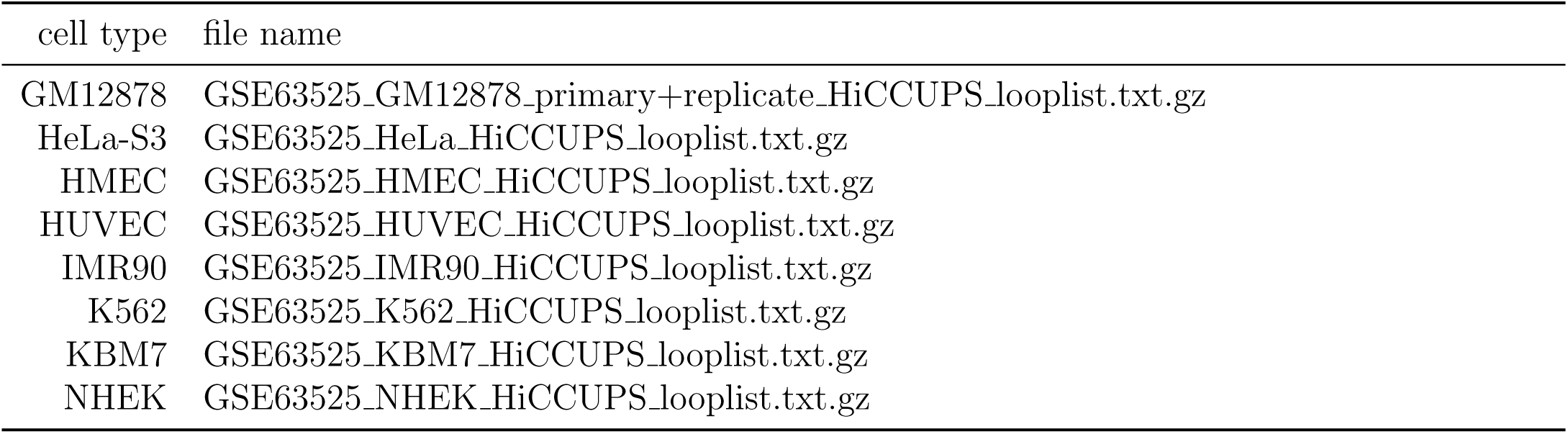
**Hi-C datasets used from GEO (accession number GSE63525).**

We used human genome assembly GRCh37/hg19 [47] for all analyses.

We used enhancers from the VISTA Enhancer Browser [28], for eye, forebrain, heart, hindbrain, limb, midbrain, and nose. We acquired the FANTOM5 blood enhancer dataset from the Promoter Enhancer Slider Selector Tool (PrESSTo) [29].

### 3D-aware genomic region enrichment

BEHST takes query regions comprising genomic loci of interest and identifies genes and annotation terms associated with these query regions through chromatin looping (Figure 2). In short, BEHST expands each query region both upstream and downstream by a query extension *e*_Q_, finds long-range interactions with one side within the expanded query region (Figure 2a–c), and then examines the distal side of these interactions (Figure 2d,i). At the distal side of a long-rage interaction, BEHST identifies *cis*-regulatory regions of protein-coding genes (Figure 2e–h). BEHST uses an upstream and downstream target extension *e*_T_ to define how far it will search for *cis*-regulatory regions. Next, BEHST creates a list of all genes in the identified *cis*-regulatory regions (Figure 2j,k). Finally, BEHST performs pathway enrichment analysis on this gene list using g:ProfileR [7] (Figure 2l,m).

We describe this procedure in more detail in the following paragraphs.

#### Extended query bounds

BEHST intersects the query regions with the long-range interaction dataset (Figure 2a,b), and then widens them by the query extension parameter *e*_Q_, in both directions. We call these widened regions the *extended query bounds*.

#### Long-range interactions

Here, we used a union of chromatin loops Hi-C datasets for the GM12878, HeLa, HMEC, HUVEC, IMR90, K562, KBM7, and NHEK cell types in Hi-C Computational Unbiased Peak Search (HiCCUPS) format [27]. We used the union of all cell types (Table 14c), rather than one specific cell type, treating the union as a repertoire of potential long-range interactions. This works in all cases, unlike requiring a mapping of a query dataset to a cell-type–specific Hi-C dataset. The appropriate Hi-C dataset to use with many queries is unclear or simply does not exist.

#### Gene annotation processing

BEHST reads a gene annotation dataset to identify potential target genes. BEHST employs APPRIS [46] to select the principal transcript for each gene. BEHST then uses the principal transcript’s identifier to extract *transcript* features from a gene annotation (Figure 2e–g). This prevents problems in downstream analysis with multiply counting genes with multiple transcripts.

#### Extended target bounds

BEHST establishes a basal *cis*-regulatory region around the principal TSS of each gene (Figure 2h). To do this, BEHST employs a strand-specific upstream and downstream adjustment (5 kbp upstream and 1 kbp downstream of the TSS). We adapted these values from GREAT [13].

From the opposite side of any chromatin loops within the extended query bounds, BEHST identifies a widened area for target search called the *extended target bounds*. For efficiency of implementation, BEHST performs this by actually extending the *cis*-regulatory regions (Figure 2i,j), but this is equivalent to extending from the target side of a chromatin loop.

#### Functional enrichment analysis

BEHST concludes by producing a list of all genes with *cis*-regulatory regions that overlap with the extended target bounds by *≥*1 bp (Figure 2k). BEHST performs functional enrichment analysis on this gene list using g:ProfileR [7], with default parameters (Figure 2l,m).

### Parameter optimization

BEHST relies on two extension parameters, the query extension *e*_Q_ and the target extension *e*_T_. To optimize these two parameters and set default values for BEHST, we performed a grid search. We ran BEHST on seven VISTA enhancer datasets (eye, forebrain, heart, hindbrain, midbrain, limb, nose) for ten values of *e*_Q_ and ten values of *e*_T_, from 100 bp to 30,000 bp. We incremented *e*_Q_ by 3000 bp, and *e*_T_ by 3100 bp in each step. We chose two different range increments to avoid identical values of the two parameters during the grid search.

For each combination of parameter values and dataset, we identified the q-value of the most significant GO term found by BEHST. We created seven matrices of most significant log_10_ (q-values), one for each of the seven VISTA datasets. Each matrix has 10 *×* 10 cells, each cell containing the most significant log_10_ (q-value) for one of 10 values of *e*_Q_ and one of 10 values of *e*_T_.

We created a 10 *×* 10 summation matrix by summing the values in all seven matrices, cell by cell. From the summation matrix, we selected the cell with the lowest total log_10_ (q-value) as parameters to use in all further analyses. This is equivalent to selecting the cell with the lowest mean. This cell corresponds to *e*_Q_ = 24,100 bp and *e*_T_ = 9400 bp.

### Negative controls

To test correctness of BEHST analyses, we created negative controls by shuffling of lists of query regions with two different procedures.

#### Total genomic random shuffle

For the total genomic random shuffle procedure, we randomly shuffled the start coordinates of each query region genome-wide, keeping region sizes identical. We performed this shuffle without regard to other genomic elements.

#### TSS-distance–preserving shuffle

The TSS-distance–preserving shuffle procedure randomly shuffles each query region across the genome, but keeps each query region as near to a TSS as it stood before shuffling. This prevents the bias inherent in a total genomic random shuffle through potentially moving query regions into a gene desert. For each query region, we calculated the distance between the region’s start and the nearest TSS of any protein-coding gene. Next, we randomly selected another TSS and moved the query region so that its start has the same distance to the new TSS as the original TSS.

### Semantic similarity of enriched terms

We used REVIGO [39] to show the similarity between the GO terms retrieved by BEHST for each query dataset. REVIGO computes semantic similarity between GO terms by considering *information content* of the terms. REVIGO defines the information content of a GO term as the negative logarithm of the frequency of that term in an annotation database. Here, we used the GO Annotations [35].

We used REVIGO’s implementation of Resnik similarity [40, 48] to estimate how much information content each pair of terms share. Resnik similarity derives from the most informative common ancestor for the two terms, and ranges in the [0*, ∞*) interval. Two terms with no informative common ancestor have Resnik similarity of 0. Terms with more informative common ancestors have higher Resnik similarities. We used Resnik similarity because it best shows correlation between gene sequence similarities and GO term similarities [49]. Resnik similarity also proves more stable than other similarity measures when used on different version of annotation databases [50, 51].

We used REVIGO analysis to exemplify differences between BEHST and GREAT on VISTA limb enhancers (Figure 5a,b). We did not perform this analysis in cases where these differences need little additional exploration. For example, GREAT retrieved only one significant GO term on the VISTA nose enhancers (Table 5. And with the FANTOM5 blood enhancers, tests with BEHST, GREAT, and ChIP-Enrich all led to expected enrichment.

## Software availability

BEHST can be used with a web browser (behst.hoffmanlab.org).

The BEHST software for Linux and macOS can be downloaded (https://bitbucket.org/hoffmanlab/behst) under the GNU General Public License version 2 (GPLv2), and can also be installed through the Bioconda [52] package distribution. We have deposited the current version of the software in Zenodo (http://doi.org/10.5281/zenodo.2174744).

## Acknowledgments

We thank Wail Ba-Alawi and the High Performance Computing team (Princess Margaret Cancer Centre) for technical assistance, and Mehran Karimzadeh (University of Toronto) and Anna Narday (University Health Network) for graphical assistance. This work was supported by the Natural Sciences and Engineering Research Council of Canada (RGPIN-2015-03948 to Michael M. Hoffman).

## Authors’ details

Davide Chicco (ORCID: 0000-0001-9655-7142) was in the Princess Margaret Cancer Centre and currently is with the Peter Munk Cardiac Centre, Toronto, Ontario, Canada.

Haixin Sarah Bi (ORCID: 0000-0001-9525-1977) was in the Princess Margaret Cancer Centre, Toronto, Ontario, Canada, and currently is in the Massachusetts Institute of Technology, Cambridge, Massachusetts, USA.

Jüri Reimand is in the Ontario Institute of Cancer Research, and in the Department of Medical Biophysics, University of Toronto, Toronto, Ontario, Canada.

Michael M. Hoffman (ORCID: 0000-0002-4517-1562) is in the Princess Margaret Cancer Centre, in the Department of Medical Biophysics and in the Department of Computer Science, University of Toronto, and in the Vector Institute, Toronto, Ontario, Canada.

Correspondence should be addressed to Michael M. Hoffman

## References

[1] Lina Wadi, Mona Meyer, Joel Weiser, Lincoln D Stein, and Jüri Reimand. Impact of outdated gene annotations on pathway enrichment analysis. Nature Methods, 13(9):705–706, 2016.

[2] Da Wei Huang, Brad T Sherman, Qina Tan, Joseph Kir, David Liu, David Bryant, Yongjian Guo, Robert Stephens, Michael W Baseler, H Clifford Lane, and Richard A Lempicki. DAVID Bioinformatics Resources: expanded annotation database and novel algorithms to better extract biology from large gene lists. Nucleic Acids Research, 35(Supplement 2):W169–W175, 2007.

[3] Mohashin Pathan, Shivakumar Keerthikumar, Ching-Seng Ang, Lahiru Gangoda, Camelia YJ Quek, Nicholas A Williamson, Dmitri Mouradov, Oliver M Sieber, Richard J Simpson, Agus Salim, Antony Bacic, Andrew F Hill, David A Stroud, Michael T Ryan, Johnson I Agbinya, John M Mariadason, Antony W Burgess, and Suresh Mathivanan. FunRich: An open access standalone functional enrichment and interaction network analysis tool. Proteomics, 15(15):2597–2601, 2015.

[4] Aravind Subramanian, Heidi Kuehn, Joshua Gould, Pablo Tamayo, and Jill P Mesirov. GSEA-P: a desktop application for Gene Set Enrichment Analysis. Bioinformatics, 23(23):3251–3253, 2007.

[5] Seth Carbon, Amelia Ireland, Christopher J Mungall, Shengqiang Shu, Brad Marshall, Suzanna Lewis, and Web Presence Working Group. AmiGO: online access to ontology and annotation data. Bioinformatics, 25(2):288–289, 2009.

[6] Edward Y Chen, Christopher M Tan, Yan Kou, Qiaonan Duan, Zichen Wang, Gabriela Vaz Meirelles, Neil R Clark, and Avi Maayan. Enrichr: interactive and collaborative HTML5 gene list enrichment analysis tool. BMC Bioinformatics, 14(1):128, 2013.

[7] Jüri Reimand, Tambet Arak, Priit Adler, Liis Kolberg, Sulev Reisberg, Hedi Peterson, and Jaak Vilo. g:Profiler–a web server for functional interpretation of gene lists (2016 update). Nucleic Acids Research, 44(W1):W83–W89, 2016.

[8] Jing Wang, Suhas Vasaikar, Zhiao Shi, Michael Greer, and Bing Zhang. WebGestalt 2017: a more comprehensive, powerful, flexible and interactive gene set enrichment analysis toolkit. Nucleic Acids Research, 45(W1):W130–W137, 2017.

[9] Cedric Simillion, Robin Liechti, Heidi EL Lischer, Vassilios Ioannidis, and Remy Bruggmann. Avoiding the pitfalls of gene set enrichment analysis with SetRank. BMC Bioinformatics, 18(1):151, 2017.

[10] Lie Li, Xinlei Wang, Guanghua Xiao, and Adi Gazdar. Integrative gene set enrichment analysis utilizing isoform-specific expression. Genetic Epidemiology, 41(6):498–510, 2017.

[11] Shijia Zhu, Tongqi Qian, Yujin Hoshida, Yuan Shen, Jing Yu, and Ke Hao. GIGSEA: genotype imputed gene set enrichment analysis using GWAS summary level data. Bioinformatics, 2018.

[12] Michael Ashburner, Catherine A Ball, Judith A Blake, David Botstein, Heather Butler, J Michael Cherry, Allan P Davis, Kara Dolinski, Selina S Dwight, Janan T Eppig, Midori A Harris, David P Hill, Laurie Issel-Tarver, Andrew Kasarskis, Suzanna Lewis, John C Matese, Joel E Richardson, Martin Ringwald, Gerald M Rubin, and Gavin Sherlock. Gene Ontology: tool for the unification of biology. Nature Genetics, 25(1):25, 2000.

[13] Cory Y McLean, Dave Bristor, Michael Hiller, Shoa L Clarke, Bruce T Schaar, Craig B Lowe, Aaron M Wenger, and Gill Bejerano. GREAT improves functional interpretation of cis-regulatory regions. Nature Biotechnology, 28(5):495–501, 2010.

[14] Ryan P Welch, Chee Lee, Paul M Imbriano, Snehal Patil, Terry E Weymouth, R Alex Smith, Laura J Scott, and Maureen A Sartor. ChIP-Enrich: gene set enrichment testing for ChIP-seq data. Nucleic Acids Research, 42(13):e105, 2014.

[15] ENCODE Project Consortium. An integrated encyclopedia of DNA elements in the human genome. Nature, 489(7414):57, 2012.

[16] Gregory P Way, Daniel W Youngstrom, Kurt D Hankenson, Casey S Greene, and Struan F Grant. Implicating candidate genes at GWAS signals by leveraging topologically associating domains. European Journal of Human Genetics, 25(11):1286, 2017.

[17] Job Dekker, Karsten Rippe, Martijn Dekker, and Nancy Kleckner. Capturing chromosome conformation. Science, 295(5558):1306–1311, 2002.

[18] Erez Lieberman-Aiden, Nynke L Van Berkum, Louise Williams, Maxim Imakaev, Tobias Ragoczy, Agnes Telling, Ido Amit, Bryan R Lajoie, Peter J Sabo, Michael O Dorschner, Richard Sandstrom, Bradley Bernstein, Michael A Bender, Mark Groudine, Andreas Gnirke, John Stamatoyannopoulos, Leonid A Mirny, Eric S Lander, and Job Dekker. Comprehensive mapping of long-range interactions reveals folding principles of the human genome. Science, 326(5950):289–293, 2009.

[19] Nynke L Van Berkum, Erez Lieberman-Aiden, Louise Williams, Maxim Imakaev, Andreas Gnirke, Leonid A Mirny, Job Dekker, and Eric S Lander. Hi-C: a method to study the three-dimensional architecture of genomes. Journal of Visualized Experiments, e1869(39), 2010.

[20] Yu Guo, Andrew A Perez, Dennis J Hazelett, Gerhard A Coetzee, Suhn Kyong Rhie, and Peggy J Farnham. CRISPR-mediated deletion of prostate cancer risk-associated CTCF loop anchors identifies repressive chromatin loops. Genome Biology, 19(1):160, Oct 2018.

[21] Scott Smemo, Juan J Tena, Kyoung-Han Kim, Eric R Gamazon, Noboru J Sakabe, Carlos Gómez-Marín, Ivy Aneas, Flavia L Credidio, Débora R Sobreira, Nora F Wasserman, Ju Hee Lee, Vijitha Puviindran, Davis Tam, Michael Shen, Joe Eun Son, Niki Alizadeh Vakili, Hoon-Ki Sung, Silvia Naranjo, Rafael D Acemel, Miguel Manzanares, Andras Nagy, Nancy J Cox, Chi-Chung Hui, Jose L Gomez-Skarmeta, and Marcelo A Nobrega. Obesity-associated variants within FTO form long-range functional connections with IRX3. Nature, 507(7492):371–375, 2014.

[22] Melina Claussnitzer, Simon N Dankel, Kyoung-Han Kim, Gerald Quon, Wouter Meuleman, Christine Haugen, Viktoria Glunk, Isabel S Sousa, Jacqueline L Beaudry, Vijitha Puviindran, Nezar A Abdennur, Jannel Liu, Per-Arne Svensson, Yi-Hsiang Hsu, Daniel J Drucker, Gunnar Mellgren, Chi-Chung Hui, Hans Hauner, and Manolis Kellis. FTO obesity variant circuitry and adipocyte browning in humans. New England Journal of Medicine, 2015(373):895–907, 2015.

[23] Laura A Lettice, Simon JH Heaney, Lorna A Purdie, Li Li, Philippe de Beer, Ben A Oostra, Debbie Goode, Greg Elgar, Robert E Hill, and Esther de Graaff. A long-range Shh enhancer regulates expression in the developing limb and fin and is associated with preaxial polydactyly. Human Molecular Genetics, 12(14):1725–1735, 2003.

[24] David U Gorkin and Bing Ren. Closing the distance on obesity culprits. Nature, 507(7492):309–310, 2014.

[25] Mijke Visser, Manfred Kayser, and Robert-Jan Palstra. HERC2 rs12913832 modulates human pigmentation by attenuating chromatin-loop formation between a long-range enhancer and the OCA2 promoter. Genome Research, 22(3):446–455, 2012.

[26] Richard C Sallari, Nicholas A Sinnott-Armstrong, Juliet D French, Ken J Kron, Jason Ho, Jason H Moore, Vuk Stambolic, Stacey L Edwards, Mathieu Lupien, and Manolis Kellis. Convergence of dispersed regulatory mutations predicts driver genes in prostate cancer. bioRxiv, page 097451, 2017.

[27] Suhas SP Rao, Miriam H Huntley, Neva C Durand, Elena K Stamenova, Ivan D Bochkov, James T Robinson, Adrian L Sanborn, Ido Machol, Arina D Omer, and Eric S Lander. A 3D map of the human genome at kilobase resolution reveals principles of chromatin looping. Cell, 159(7):1665–1680, 2014.

[28] Axel Visel, Simon Minovitsky, Inna Dubchak, and Len A Pennacchio. VISTA Enhancer Browser–a database of tissue-specific human enhancers. Nucleic Acids Research, 35(Supplement 1):D88–D92, 2007.

[29] Robin Andersson, Claudia Gebhard, Irene Miguel-Escalada, Ilka Hoof, Jette Bornholdt, Mette Boyd, Yun Chen, Xiaobei Zhao, Christian Schmidl, Takahiro Suzuki, Evgenia Ntini, Erik Arner, Eivind Valen, Kang Li, Lucia Schwarzfischer, Dagmar Glatz, Johanna Raithel, Berit Lilje, Nicolas Rapin, Frederik Otzen Bagger, Mette Jorgensen, Peter Refsing Andersen, Nicolas Bertin, Owen Rackham, A Maxwell Burroughs, J Kenneth Baillie, Yuri Ishizu, Yuri Shimizu, Erina Furuhata, Shiori Maeda, Yutaka Negishi, Christopher J Mungall, Terrence F Meehan, Timo Lassmann, Masayoshi Itoh, Hideya Kawaji, Naoto Kondo, Jun Kawai, Andreas Lennartsson, Carsten O Daub, Peter Heutink, David A Hume, Torben Heick Jensen, Harukazu Suzuki, Yoshihide Hayashizaki, Ferenc Muller, Alistair R Forrest, Piero Carninci, Michael Rehli, and Sandelin Albin. An atlas of active enhancers across human cell types and tissues. Nature, 507(7493):455–461, 2014.

[30] Len A Pennacchio, Nadav Ahituv, Alan M Moses, Shyam Prabhakar, Marcelo A Nobrega, Malak Shoukry, Simon Minovitsky, Inna Dubchak, Amy Holt, Keith D Lewis, Ingrid Plajzer-Frick, Jennifer Akiyama, Sarah De Val, Veena Afzal, Brian L Black, Olivier Couronne, Michael B Eisen, Visel Axel, and Edward M Rubin. In vivo enhancer analysis of human conserved non-coding sequences. Nature, 444(7118):499, 2006.

[31] Diogo S Castro, Ben Martynoga, Carlos Parras, Vidya Ramesh, Emilie Pacary, Caroline Johnston, Daniela Drechsel, Mélanie Lebel-Potter, Laura Galinanes Garcia, Charles Hunt, Dirk Dolle, Angela Bithell, Laurence Ettwiller, Noel Buckley, and Franois Guillemot. A novel function of the proneural factor ASCL1 in progenitor proliferation identified by genome-wide characterization of its targets. Genes & Development, 25(9):930–945, 2011.

[32] Rimantas Kodzius, Miki Kojima, Hiromi Nishiyori, Mari Nakamura, Shiro Fukuda, Michihira Tagami, Daisuke Sasaki, Kengo Imamura, Chikatoshi Kai, Matthias Harbers, Yoshihide Hayashizaki, and Piero Carninci. CAGE: cap analysis of gene expression. Nature Methods, 3(3):211, 2006.

[33] Jüri Reimand, Meelis Kull, Hedi Peterson, Jaanus Hansen, and Jaak Vilo. g:Profiler–a web-based toolset for functional profiling of gene lists from large-scale experiments. Nucleic Acids Research, 35(Supplement 2):W193–W200, 2007.

[34] g:Profiler. g:Profiler help. https://biit.cs.ut.ee/gprofiler/help.cgi7help_id=5, 2016 (accessed on 6 July 2018).

[35] Gene Ontology Consortium. Gene Ontology annotations and resources. Nucleic Acids Research, 41(D1):D530–D535, 2013.

[36] Bronwen L Aken, Sarah Ayling, Daniel Barrell, Laura Clarke, Valery Curwen, Susan Fairley, Julio Fernandez Banet, Konstantinos Billis, Carlos García Girón, Thibaut Hourlier, Kevin Howe, Andreas Kahari, Felix Kokocinski, Fergal J Martin, Daniel N Murphy, Rishi Nag, Magali Ruffier, Michael Schuster, Y Amy Tang, Jan-Hinnerk Vogel, Simon White, Amonida Zadissa, Paul Flicek, and Stephen M Searle. The Ensembl gene annotation system. Database, 2016(Database update):1–19, 2016.

[37] Robert C Gentleman, Vincent J Carey, Douglas M Bates, Ben Bolstad, Marcel Dettling, Sandrine Dudoit, Byron Ellis, Laurent Gautier, Yongchao Ge, Jeff Gentry, Kurt Hornik, Torsten Hothorn, Wolfgang Huber, Stefano Iacus, Rafael Irizarry, Friedrich Leisch, Cheng Li, Martin Maechler, Anthony J Rossini, Gunther Sawitzki, Colin Smith, Gordon Smyth, Luke Tierney, Jean YH Yang, and Jianhua Zhang. Bioconductor: open software development for computational biology and bioinformatics. Genome Biology, 5(10):R80, 2004.

[38] Ryan P Welch, Chee Lee, Paul M Imbriano, Snehal Patil, Terry E Weymouth, R Alex Smith, Laura J Scott, and Maureen A Sartor. ChIP-Enrich - gene set enrichment testing for ChIP-seq data and other sets of genomic regions. http://chip-enrich.med.umich.edu/, 2013 (accessed on 16 December 2018).

[39] Fran Supek, Matko Bošnjak, Nives Skunca, and Tomislav Smuc. REVIGO summarizes and visualizes long lists of Gene Ontology terms. PLOS ONE, 6(7):e21800, 2011.

[40] Philip Resnik. Using information content to evaluate semantic similarity. In Proceedings of IJCAI’95 – the 14th International Joint Conference on Artificial Intelligence, pages 448–453, 1995.

[41] Catia Pesquita, Daniel Faria, Andre O Falcao, Phillip Lord, and Francisco M Couto. Semantic similarity in biomedical ontologies. PLOS Computational Biology, 5(7):e1000443, 2009.

[42] Daniel Barrell, Emily Dimmer, Rachael P Huntley, David Binns, Claire ODonovan, and Rolf Apweiler. The GOA database in 2009 - an integrated Gene Ontology Annotation resource. Nucleic Acids Research, 37(supplement 1):D396–D403, 2008.

[43] Trevor F Cox and Michael A Cox. Multidimensional scaling. Chapman and Hall/CRC, 2000.

[44] Fran Supek, Matko Bošnjak, Nives Skunca, and Tomislav Šmue. REVIGO summarizes and visualizes long lists of Gene Ontology terms. http://revigo.irb.hr/, 2017 (accessed on 15 January 2019).

[45] Thomas Derrien, Rory Johnson, Giovanni Bussotti, Andrea Tanzer, Sarah Djebali, Hagen Tilgner, Gregory Guernec, David Martin, Angelika Merkel, David G Knowles, Julien Lagarde, Lavanya Veeravalli, Xiaoan Ruan, Yijun Ruan, Timo Lassmann, Piero Carninci, James B Brown, Leonard Lipovich, Jose M Gonzalez, Mark Thomas, Carrie A Davis, Ramin Shiekhattar, Thomas R Gingeras, Tim J Hubbard, Cedric Notredame, Jennifer Harrow, and Roderic Guigo. The GENCODE v7 catalog of human long noncoding RNAs: analysis of their gene structure, evolution, and expression. Genome Research, 22(9):1775–1789, 2012.

[46] Jose Manuel Rodriguez, Paolo Maietta, Iakes Ezkurdia, Alessandro Pietrelli, Jan-Jaap Wes-selink, Gonzalo Lopez, Alfonso Valencia, and Michael L Tress. APPRIS: annotation of principal and alternative splice isoforms. Nucleic Acids Research, 41(D1):D110–D117, 2012.

[47] Valerie Schneider and Deanna Church. Genome Reference Consortium. National Center for Biotechnology Information, 2013.

[48] Philip Resnik. Semantic similarity in a taxonomy: an information-based measure and its application to problems of ambiguity in natural language. Journal of Artificial Intelligence Research, 11:95–130, 1999.

[49] Xiang Guo, Rongxiang Liu, Craig D Shriver, Hai Hu, and Michael N Liebman. Assessing semantic similarity measures for the characterization of human regulatory pathways. Bioinformatics, 22(8):967–973, 2006.

[50] Norberto Díaz-Díaz and Jesús S Aguilar-Ruiz. GO-based functional dissimilarity of gene sets. BMC Bioinformatics, 12(1):360, 2011.

[51] Davide Chicco, Fernando Palluzzi, and Marco Masseroli. Novelty indicator for enhanced prioritization of predicted Gene Ontology annotations. IEEE/ACM Transactions on Computational Biology and Bioinformatics, 15(3):954–965, 2018.

[52] Björn Grüning, Ryan Dale, Andreas Sjödin, Jillian Rowe, Brad A Chapman, Christopher H Tomkins-Tinch, Renan Valieris, The Bioconda Team, and Johannes Köster. Bioconda: sustainable and comprehensive software distribution for the life sciences. Nature Methods, 15(7):475–476, 2018.

